# Human whole-epigenome modelling for clinical applications with Pleiades

**DOI:** 10.1101/2025.07.16.665231

**Authors:** Pouya Niki, Christoforos Nalmpantis, Javkhlan-Ochir Ganbat, Donal Byrne, Pooja Kathail, Ching Fang, Alfred Wong, Andrey Karailiev, Husam Babikir, Anjeet Jhutty, Luca Giacomoni, Francisco M Marín-Zamora, Arihant Kamdar, Nicholas K. Wang, Will Rowe, Timing Liu, Netanel Loyfer, Robert Sugar, Augustinas Malinauskas, Khaled Saab, Mark Bissell, Dron Hazra, Michael T. Pearce, Archa Jain, Daniel Balsam, Sofia Toniolo, Masud Husain, Sanjay G. Manohar, Sian A. Thompson, Henrik Zetterberg, Hannah Madan, Ivan Koychev, Jonathan C. M. Wan, Ravi Solanki

**Author notes:** These authors contributed equally to this work (Pouya Niki, Christoforos Nalmpantis, Javkhlan-Ochir Ganbat, Donal Byrne, Pooja Kathail).

## Abstract

Gene regulation in humans extends beyond the four-letter genetic code. DNA methylation, in particular, functions as a critical epigenetic regulator, dynamically programming cellular identity, adapting gene expression in response to environmental cues, and underpinning the onset and progression of numerous diseases. Here we present Pleiades, a series of whole-genome epigenetic foundation models spanning three sizes: 90M, 600M, and 7B parameters. Pleiades is trained upon an extensive proprietary corpus of human methylation and genomic data, totalling 1.9T tokens. We introduce alignment embeddings and stacked hierarchical attention techniques to provide precise epigenetic modelling without the need for extended context lengths. Collectively, these advances enable Pleiades to perform a diverse range of down-stream biological and clinical tasks, including genomic regulatory prediction, realistic generation of cell-free DNA fragments and fragment-level cell-type-of-origin classification, within a unified and scalable computational framework. We specifically apply Pleiades to the early detection of clinical Alzheimer’s disease and Parkinson’s disease from plasma cell-free DNA, achieving high-accuracy detection (AUROC 0.89 for AD and 0.84 for PD) using a minimally invasive blood test. Combined with plasma pTau-217, Pleiades reaches an AUROC of 0.97 for AD, underscoring the promise of multimodal epigenomic and proteomic approaches. Using mechanistic interpretability, we ground Pleiades’ latent features in interpretable biological signals, relating its Alzheimer’s predictions to fragmentomic and epigenomic properties of cfDNA. These findings demonstrate the potential of genome-wide epigenomic modelling as a clinically translatable paradigm for diagnostics and precision medicine, though prospective validation in larger, more diverse cohorts will be required to confirm clinical utility.

## 1 Introduction

The creation of tools to understand human biology has been critical for the advancement of health and the prevention of disease for centuries [1–9]. Applications of artificial intelligence and advanced language modelling to the life sciences have established a new era for the discovery of biological knowledge, promising diagnostics and therapeutics across a range of complex human diseases [10–14].

Many such tools and models have focused upon the human genome, effectively capturing the underlying statistics of DNA patterns and coevolutions [11, 15–19]. However, these models do not capture the epigenome, the set of dynamic environmental and chemical changes to the genetic code critical for organismal development, cellular fate, and both the onset and progression of multiple diseases [20–25]. Of the variety of epigenetic alterations, DNA methylation is a critical class [26, 27]. Its study has led to the development of diagnostics for cancer detection and identification of novel mechanisms responsible for age-related diseases [28–31].

The dynamic role of DNA methylation is increasingly evident in the pathophysiology of neurodegenerative conditions, including Alzheimer’s disease (AD), Parkinson’s disease (PD), and amyotrophic lateral sclerosis (ALS) [32–37]. Dementia, which frequently arises from such neurodegeneration, is among the leading causes of morbidity and mortality in the world: first in the UK, sixth in the USA, and seventh globally [38–40]. It has historically been challenging to study due to the lack of ability to study pathology in living patients, poor translation of animal models, heterogeneity of disease, and an insidious disease onset with pathology preceding symptoms by up to two decades [41–44]. Recent cfDNA-based studies have demonstrated altered fragment profiles and methylation signatures in AD and PD patient plasma [45–47], suggesting that cfDNA may serve as a non-invasive window into central nervous system pathology, with granularity that may allow quantification of disease heterogeneity at an individual level. The lack of precision diagnostic tools poses a challenge to both patient outcomes and clinical trials for these neurodegenerative conditions [48–50]. While recent advances in proteomic tests for AD (most notably plasma pTau-217) and PD (mainly cerebrospinal fluid tests detecting pathological alpha-synuclein seeds) show promise, each interrogates a single pathology with limited appreciation of co-pathologies, and detection of AD/PD pathology does not necessarily associate with clinical conversion, leaving them limited to a subset of patients and disease stages, and leading to real-world clinical ambiguities [51–56].

In recent years, similar challenges faced in oncology have been addressed with the study and application of cell-free DNA (cfDNA), fragments of DNA that freely circulate in a variety of biofluids such as plasma and cerebrospinal fluid (CSF) [28, 57]. Importantly, the methylation status of cfDNA is indicative of its cellular origin as well as of changes that may occur throughout the disease process [57, 58]. To date, cfDNA has been successfully utilised for the early detection of cancers, monitoring of treatment response, and the discovery of novel biology for downstream therapeutic application [57, 59–63].

Here we present Pleiades, a series of biological foundation models for the human epigenome at three scales — 90M, 600M, and 7B parameters. Pleiades couples an autoregressive transformer decoder with *alignment embeddings* — single-base positional embeddings that locate every nucleotide within the human genome — and a multi-tier *hierarchical attention transformer* that pools individual cfDNA fragments into sample-level representations. The models jointly capture DNA sequence and methylation, trained on a 1.9T-token corpus we curated from a high-quality tissue-specific methylation atlas, plasma-derived cfDNA, and a graph of human genomic diversity [58, 64, 65].

Pleiades is a *generalist* epigenomic foundation model: from a single set of pretrained weights, it rivals or outperforms task-specific specialist models across a wide range of biological tasks. On the Nucleotide Transformer benchmark of human genomic annotations, Pleiades achieves state-of-the-art performance, with even the smallest 90M model outperforming DNA-only baselines many times its size [12, 15]. The frozen representations of Pleiades 7B further surpass recent genomic foundation models, including the long-context Evo 2 [19] and AlphaGenome [66], on histone-mark prediction without any task-specific fine-tuning. Pleiades 7B also generates plasma cfDNA fragments *in silico* with high fidelity, recapitulating nucleosome-driven fragment-length distributions *de novo*. Plasma cfDNA is dominated by hematopoietic fragments, while disease-relevant signal often originates from rare cell types such as neurons and microglia; resolving cellular origin at the per-fragment level is therefore critical for sensitive diagnostic modelling. Pleiades classifies the cellular origin of individual cfDNA fragments, performing on par with established sample-level deconvolution tools such as UXM and CelFiE while enabling enrichment of plasma samples for rare, disease-relevant cell types.

We then apply Pleiades to detection of clinical neurodegenerative disease from plasma cfDNA in two independent real-world cohorts spanning Alzheimer’s disease (AD) and Parkinson’s disease (PD) [45, 46, 67, 68]. Pleiades discriminates patients from age- and sex-matched controls in both conditions, outperforming established cfDNA baselines in each cohort. Combined with plasma pTau-217, Pleiades reaches an AUROC of 0.97 for AD, suggesting multi-modal integration of cfDNA and proteomic biomarkers as a promising direction for future work. Mechanistic interpretability via sparse autoencoders grounds the AD classifier’s latent features in interpretable biological signals, relating its predictions to fragmentomic and epigenomic properties of cfDNA, offering a candidate hypothesis for cfDNA biomarker design in neurodegenerative disease. Together, these results provide a proof of concept for a generalisable framework for blood-based biomarker discovery, with potential to extend to many other diseases.

## 2 Results

### 2.1 Pleiades

The Pleiades series scales to 7B parameters and is trained upon a unique data corpus of 1.9T tokens of human methylation and genomic data, using a context length of 1,024 tokens (Fig. 1a).

**Fig. 1:**
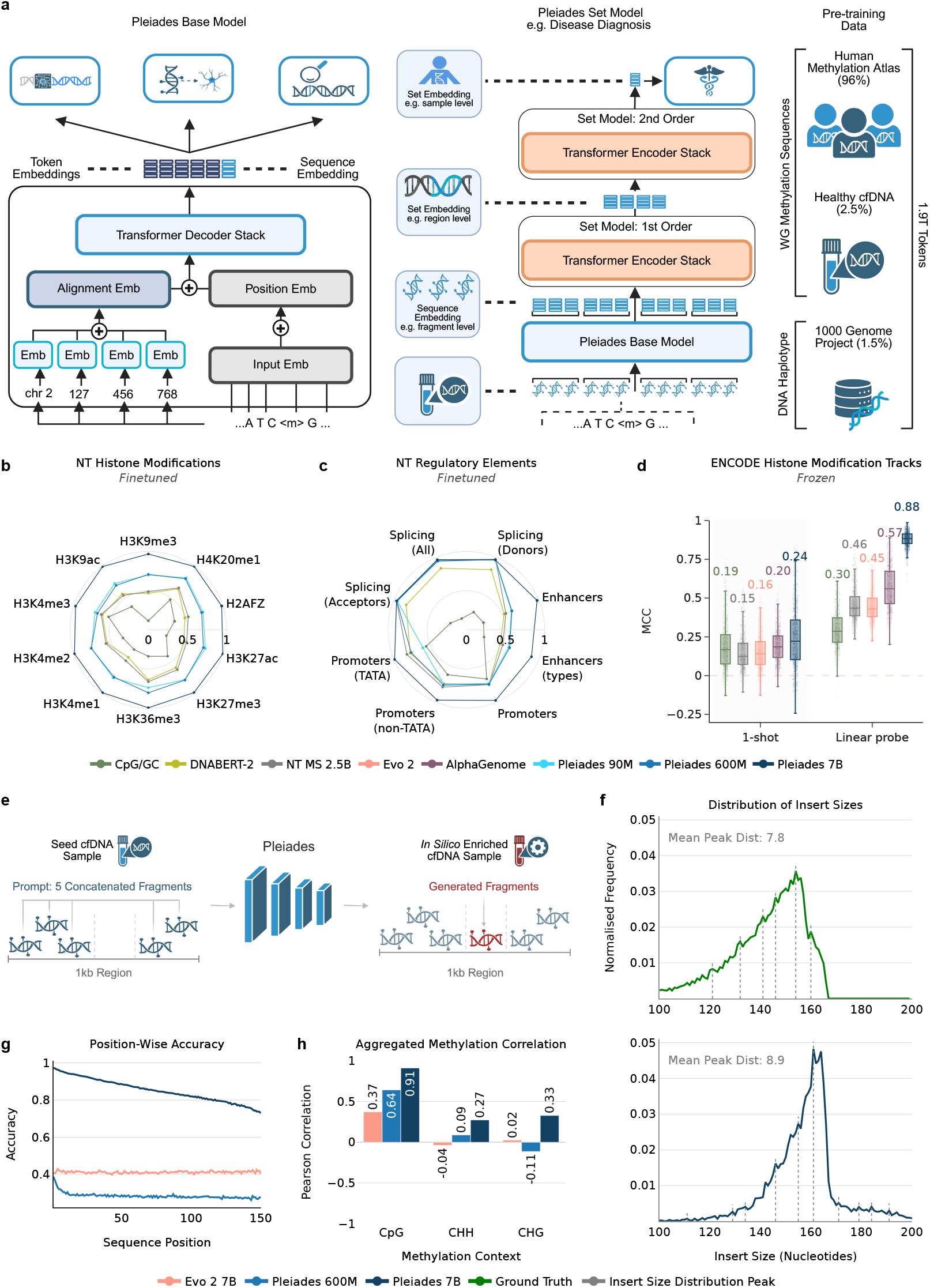
Epigenomic Foundation Modelling with Pleiades. **(a)** *Base model* : token-level inputs (sequence, methylation, position, alignment embeddings) are processed by a transformer decoder to per-token and sequence-level embeddings, used for sequence-level tasks. *Hierarchical set model* : fragment-level embeddings are aggregated within genomic windows into region-level vectors, then pooled by a second transformer encoder into a sample-level embedding for set- and sample-level tasks (e.g. disease diagnosis). **(b)** Fine-tuned Matthews correlation coefficient (MCC) on Nucleotide Transformer (NT) Benchmark histone-modification tasks (per mark), against NT MS 2.5B, DNA-BERT2, and a CpG/GC baseline (non-learned: CpG ratio and GC content). **(c)** Finetuned MCC on NT gene-regulatory-element tasks (enhancer, promoter, splice site) against the same baselines. **(d)** Frozen-representation MCC on ENCODE histone-modification tracks under 1-shot and linear-probe regimes, comparing Pleiades 7B against CpG/GC, NT MS 2.5B, Evo 2, and AlphaGenome. Pleiades 7B significantly outperforms every baseline in both regimes (two-sided paired *t*-test across tracks, Benjamini–Hochberg corrected, *p*_BH_ *<* 10^−3^). Panels (b)–(d) share a legend; only the models named per panel are shown. **(e)** cfDNA generation: five observed cfDNA fragments prompt the model to generate a sixth, non-overlapping fragment. **(f)** Insert-size distribution of generated fragments versus ground truth; dashed line marks the distribution peak. **(g)** Position-wise nucleotide accuracy along the generated fragment. **(h)** Pearson correlation between generated and ground-truth methylation by sequence context (CpG, CHH, CHG).

#### 2.1.1 Pretraining

Pleiades adopts an autoregressive transformer decoder architecture, provided at three scales — 90M, 600M, and 7B parameters. To effectively pretrain Pleiades, we required large amounts of high-quality sequence data from a variety of human cell types and samples. Existing public resources such as the NIH Roadmap Epigenomics Mapping Consortium and ENCODE Consortium are extensive, but heterogeneous sample quality limits their utility for foundation modelling [22, 58, 69]. To overcome this, we curated our own consolidated corpus of human methylation and genomic data:

1. **DNA methylation atlas of normal human cell types**: whole-genome bisulfite sequencing (WGBS) of 39 cell-type groups obtained via fluorescence-activated cell sorting from 205 tissue samples across 137 healthy donors [58].
2. **Plasma-derived cfDNA**: WGBS and enzymatic methylation sequencing (EM-Seq) of plasma cell-free DNA from 20 healthy individuals [64].
3. **Human genome diversity**: a genome graph representative of the 1000 Genomes Project [65].

Comprehensive dataset preparation and tokenisation details are provided in Section 4.1.4.

#### 2.1.2 Alignment Embeddings

Capturing long-range regulatory interactions, which can span megabases, requires representing precise genomic context within fixed-length transformer inputs [70]. To this end, we introduce *alignment embeddings* (AEs), which encode each nucleotide’s absolute genomic coordinate directly into the model’s sequence representation (Methods, Section 4.1.2). By embedding positional information, AEs equip Pleiades to recognise subtle yet biologically meaningful differences among genomic and methylomic sequences, without relying upon prohibitively large transformer context windows.

For each nucleotide position within a read, we decompose its GRCh38 genomic position (chromosome:position) into four integers: chromosome, millions position, thousands position and ones position. Each of the four integers passed through its own embedding table to produce four vectors, which were concatenated to form the Alignment Embedding token. These embeddings were trained jointly with the model, covering every base, thereby capturing long-range epigenomic structure inside a fixed-length transformer.

Unlike recent positional embedding approaches such as CpGPT [16] that encode location only for CpG dinucleotides (≈ 30 million loci in GRCh38), our AEs encode the absolute chromosome and single-base offset for every nucleotide in the human genome (≈ 3.1 billion positions; Fig. 1a). Full details of the AEs are outlined in Methods, Section 4.1.2.

#### 2.1.3 Hierarchical Set Modelling

Many downstream clinical-genomics tasks require reasoning over an entire biological sample rather than individual reads. For such tasks, we treat each sample’s cfDNA reads as a set of sequences — typically 10^8^–10^9^ in size — and require an aggregation mechanism that scales to this regime.

To summarise these large sets, we adopt a multi-tier *Hierarchical Attention Transformer* (HAT), inspired by Chalkidis *et al*. [71]. In our implementation, we stack *N* HAT blocks: each block pools fixed-size groups of lower-level tokens and passes the condensed representation upward such that successively higher-order summaries are built in *N* steps. The top-level token yields a compact sample-level embedding that feeds directly into a classifier. This hierarchical pooling preserves long-range dependencies, allowing Pleiades to tackle large sequencing tasks without the need for long context methods or alternative base architectures. Full architectural and training details appear in Section 4.1.3.

### 2.2 DNA Sequence Classification Benchmarks

To systematically evaluate the capabilities of the Pleiades architecture and epigenetic-focused pretraining, we benchmarked the model on established genomic classification tasks originally introduced by Nucleotide Transformer [15]. These benchmarks include the ability of a model to identify a variety of genomic features, such as promoters, enhancers, splice sites, and histone modification sites, and are commonly used to assess the predictive performance of DNA sequence models [12, 15].

During our analysis, we identified a positional bias in the original Nucleotide Transformer dataset: negative sequences consistently begin at genomic positions divisible by 1,000, providing a trivial cue for distinguishing positive from negative examples and inflating reported performance (Supp. Fig. S2). To address this, we introduced a randomised − jitter in the range [− 500, 499] to the start positions of negative sequences, effectively removing the positional bias from our findings (Supplementary Fig. S2). We refer to this as the Unbiased Nucleotide Transformer Benchmark. A comprehensive explanation of this adjustment is detailed in Supplementary Section S1.

We benchmarked Pleiades models (90M, 600M, and 7B) against two popular baseline genomic foundation models: the largest Nucleotide Transformer model (Multi-species, 2.5B parameters; NT MS 2.5B) [15] and DNA-BERT2 [12]. We also included a non-learned CpG/GC composition baseline (CpG ratio and GC content). Each model was fine-tuned for exactly five epochs on our Unbiased Nucleotide Transformer Benchmark, whereas the CpG/GC baseline used these fixed sequence-composition features directly. To alleviate potential distribution shift effects between predominantly methylomic pretraining data for Pleiades (98.5% of total pretraining data) and this purely genomic fine-tuning dataset, small Pleiades models (90M, 600M) were fine-tuned for one epoch on the DNA sub-portion of the pretraining dataset prior to evaluation (1.5% of total pretraining data). Pleiades 7B did not undergo any specific DNA fine-tuning.

Pleiades 7B achieves the highest Matthews correlation coefficient (MCC) on 15 of 18 tasks and is on par with baselines on the remaining three, with a macro-average MCC of 0.98 (Fig. 1b,c; Supp. Fig. S1) (Methods, Section 4.2). The smaller 90M and 600M models exceed baselines on 12 of 18 tasks, with macro-average MCCs of 0.76 and 0.77 — well above the 0.63 of DNABERT-2 and 0.67 of NT MS 2.5B. Full results are detailed in Table S1.

On histone modification prediction tasks, even small Pleiades models outperformed DNA-only baselines (Fig. 1b). Pleiades 90M outperformed the strongest baseline (NT MS 2.5B) despite carrying 27× fewer parameters. This highlights the power of DNA methylation to better capture genome regulatory information than pure DNA models [72, 73].

To test whether this advantage arises from Pleiades exploiting CpG density as a proxy for methylation, we stratified prediction errors by per-locus CpG content across the 9 positively-aligned histone-mark tasks (Supp. Table S2). DNABERT-2 and NT MS 2.5B exhibit strongly CpG-conditioned errors — a ∼3× higher false-positive rate at CpG-rich loci and a ∼4× higher false-negative rate at CpG-poor loci — whereas Pleiades 7B shows essentially none, indicating that methylation-aware pre-training reduces rather than exploits reliance on CpG content as a shortcut for histone mark prediction.

We further investigate the Pleiades series by examining the few-shot learning capabilities on these tasks. Supp. Fig. S1c shows the performance on the representative histone modification task for H3K27ac for all models under study, trained for one epoch on the entire dataset. Pleiades 7B achieves near-perfect MCC (0.9925) after training on only 152 examples. No other model exhibits comparable few-shot capability; baselines achieve lower performance even after training on the full ∼30,000-sample dataset for one epoch.

Beyond full fine-tuning, we tested whether the representations themselves are informative without any task-specific training. We extracted frozen embeddings from Pleiades 7B and the NT MS 2.5B, Evo 2, and AlphaGenome baselines, and added a non-learned CpG/GC composition control (2-dimensional: CpG ratio and GC content). Each representation was scored through an identical pipeline under two regimes: a 1-shot nearest-prototype classifier and a linear probe trained on the full task labels (Methods, Section 4.2.1). All models were evaluated on the same 1 kb window as Pleiades, except AlphaGenome, whose fixed input requires a minimum of 2 kb. Among these models, only AlphaGenome was pretrained directly on genomic tracks of this type. On 884 ENCODE histone-modification tracks, the linear probe gave Pleiades 7B a median MCC of 0.88, ahead of AlphaGenome (2 kb, 0.56), NT MS 2.5B (0.44), Evo 2 (1 kb, 0.43), and CpG/GC (0.29); Pleiades 7B ranked first on all 884 tracks (two-sided paired *t*-test across tracks, Benjamini–Hochberg corrected, *p*_BH_ *<* 10^−3^ against each baseline), and this ordering held when tracks were aggregated to ten canonical histone marks (Fig. 1d; Supp. Fig. S1g). The 1-shot regime followed the same ranking, with Pleiades 7B at a median MCC of 0.22, above every baseline shown (*p*_BH_ *<* 10^−3^).

The same evaluation on the NT benchmark tasks placed Pleiades 7B first on every task under the linear probe, at median MCCs of 0.99 (10 histone tasks) and 0.96 (8 regulatory tasks) (*p*_BH_ ≤ 2 × 10^−4^); under the 1-shot regime it led on the histone tasks (*p*_BH_ ≤ 3 × 10^−4^), while differences on the regulatory tasks were not significant (Supp. Fig. S1d,e). Because Evo 2 and AlphaGenome are built to read longer sequence context than Pleiades, we additionally evaluated them at 8 kb to test whether more context narrows the gap. On ENCODE, the longer context raised their linear-probe medians to 0.50 (Evo 2) and 0.64 (AlphaGenome), both still below Pleiades 7B at 1 kb (*p*_BH_ *<* × 10^−3^); in the 1-shot regime, AlphaGenome at 8 kb exceeded Pleiades 7B by a median of 0.017 MCC (*p*_BH_ ≈ 10^−3^) (Supp. Fig. S1f).

### 2.3 Epigenomic Sequence Generation

We next assessed Pleiades’ generative capabilities on plasma cfDNA, evaluating its ability to produce realistic *in silico* biological sequences (Fig. 1e).

We frame the task on cfDNA bio-samples, held out during pretraining, each sequenced to a depth of 20 − 50*x*. In each sample, we randomly split fragments into a 10% seed set (used to construct prompts) and a 90% ground-truth set (used as generation targets). Each prompt comprised five full seed-set fragments from the same 1kb region followed by the first three nucleotides of a non-overlapping ground-truth fragment from the same region. We focused on 68 repeat-masked high-coverage 1kb regions. A full list of these regions can be found in Table S3.

We established three classes of evaluation metrics to assess generation quality at three complementary resolutions:

i. *Nucleotide fidelity* : Per-position accuracy (Fig. 1g) and longest-common-subsequence (LCS) length relative to ground truth fragment length (Fig. S3b).
ii. *Methylome concordance*: Pearson correlation of aggregated methylation ratios (CpG/CHG/CHH) across 1kb bins (Fig. 1h) and analysis of methylation with respect to cytosine context (Fig. S3c).
iii. *Fragmentomics*: Insert-length distribution and insert size distribution peak periodicity (Fig. 1f; Fig. S3a).

For baseline comparisons, we chose Evo 2 7B, a DNA-only language model trained upon genomic sequences across multiple species [19]. This allowed for a direct assessment of the utility of DNA-only models without specialised training or data inputs to capture complex epigenetic signatures.

Assessing nucleotide accuracy, Pleiades 7B achieved 97% accuracy of the first nucleotide and 73% at base 150, for a mean of 83% (Fig. 1g). Pleiades 600M started at 40% and declined to 19%, for a mean of 25%. In comparison, Evo 2 7B remained largely static across the trace (mean 42%). We observed that parameter scaling substantially improved single-nucleotide level accuracy.

LCS analysis indicated similar results (Fig. S3b). Pleiades 7B reproduced *on average* 85% of ground-truth fragments contiguously (≈ 125 nucleotides of a 150 nucleotide window). The model achieved a *median* relative LCS length of 0.98. This indicates that using the 10% seed set alone, 50% of generated fragments matched at least 98% of the original sequence. In contrast, Pleiades 600M and Evo 2 7B had a relative LCS length of only 0.07 and 0.08 (≈ 11–12 nt) on average, respectively.

We next assessed the concordance of methylation across cytosine context (Fig. S3c). Methylation can occur across three genomic contexts: CpG, CHG, and CHH. While the vast majority of methylation in the human genome occurs at CpG sites, non-CpG methylation is especially critical for brain biology and brain disease [32, 74–76]. In our ground truth set, approximately 91% of all methylation occurs at CpG sites, with only 7% at CHH and 2% at CHG sites. Methylation–context ratio analysis shows that Evo 2 7B under-represented CpG contexts by 24.1% while over-generating CHH by 24.8%, indicating a bias towards non-canonical methylation sites. In contrast, Pleiades 600M reduced these errors to +8.2% (CpG) and −6.6% (CHH) and Pleiades 7B further to +4.0% (CpG) and −3.1% (CHH). All three models maintained CHG deviations below 1.6% (Figure S3c).

We then compared aggregated methylation correlation between predicted and true across context (Fig. 1h). Pleiades 7B attains a Pearson correlation of 0.91 with ground-truth CpG methylation, in contrast to 0.64 for Pleiades 600M and 0.37 for Evo 2 7B. Correlations for non-CpG methylation are lower, with the best performance seen with Pleiades 7B in each of CHH (0.27 vs. 0.09 Pleiades 600M and −0.04 Evo 2 7B) and CHG (0.33 vs. −0.11 600M and 0.02 Evo 2 7B). As expected, larger scale and methylation-focussed pretraining together reduce context-distribution biases and improve locus-specific methylation recall.

*In silico* generated fragments were plotted by insert size (Fig. 1f; Fig. S3a). The distribution reflects the canonical nucleosome-related architecture with a modest right-shift in their length distribution. Modal length increased from 154nt in the ground-truth library to 163nt in the generated set (Δ = +9 nt; +5.8%). Rotational phasing of cfDNA, quantified by the spacing between successive mini-peaks in the insert size spectrum, was virtually preserved (8.9 ± 0.3nt *in silico* vs. 7.8 ± 0.2nt empirically). Interestingly, these higher-order chromatin signatures arise *de novo*; the model was never *directly* supplied with fragment-length targets or nucleosome annotations, underscoring that base-resolution sequence pretraining alone suffices to learn nucleosome organisation.

Collectively, these results show that scaling from 600M to 7B parameters raises mean per-nucleotide accuracy from 25% to 83%, increases relative LCS from 0.07 to 0.85, sharpens CpG methylation correlation from 0.64 to 0.91, and preserves nucleosome-driven fragment lengths. In contrast, the DNA-only baseline underperforms across all metrics despite matching the 7B parameter budget.

### 2.4 Cell Type-of-Origin (CToO)

Cell Type-of-Origin (CToO) refers to the specific cell type from which a circulating cfDNA fragment originates. As plasma-derived cfDNA inherently comprises fragments from diverse cell types, accurate determination of the CToO of cfDNA fragments is necessary for precise evaluation of a specific tissue’s physiological state and facilitates early diagnosis, disease monitoring, and targeted therapeutic interventions [58, 62]. In neurodegenerative disease, this capability is particularly relevant: disease-relevant cfDNA fragments originating from neurons, microglia, or oligodendrocytes constitute a small fraction of total plasma cfDNA, which is dominated by haematopoietic contributions [58]. Fragment-level CToO classification enables two clinically meaningful capabilities beyond standard deconvolution: (i) enrichment of rare disease-relevant fragments prior to diagnostic modelling, potentially improving signal-to-noise for early detection; and (ii) per-fragment attribution of cell-type identity for longitudinal tissue damage monitoring (e.g., tracking neuronal cfDNA fraction changes following therapeutic intervention).

In contrast, existing cell type deconvolution methods typically produce aggregate, sample-level estimates, rather than per fragment predictions [58, 64]. Here, we propose the Cell Type-of-Origin task, an approach that leverages methylation patterns of individual cfDNA fragments (100–300 bp) to determine their originating cell type on a per fragment level (Fig. 2a). The hierarchical architecture of Pleiades, which operates at the fragment level before aggregating to sample-level predictions, is specifically designed to exploit these fine-grained classifications, a capability not available to methods that operate only on aggregated methylation summaries.

**Fig. 2:**
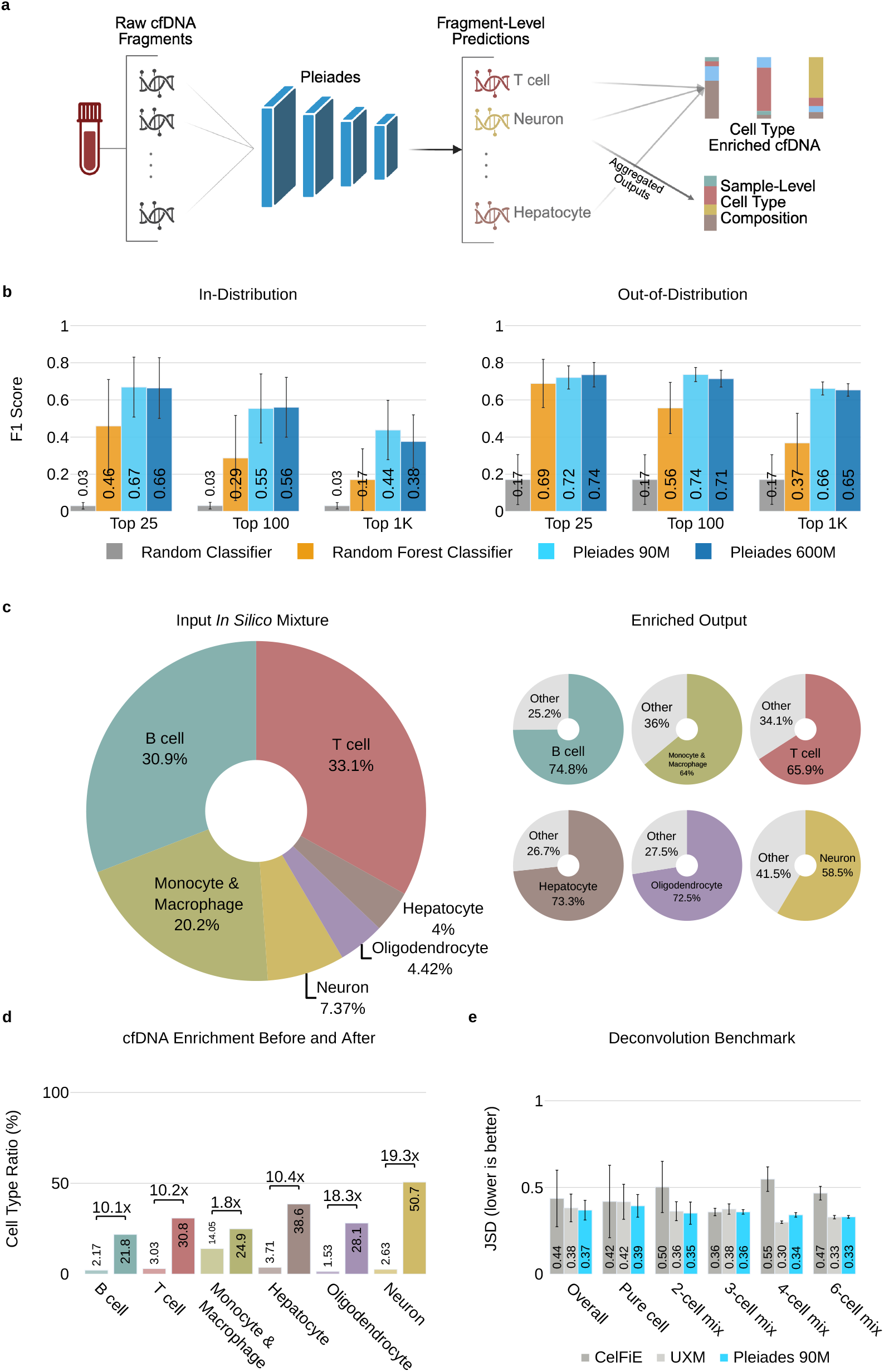
Pleiades Predicts the Cell Type-of-Origin of Fragments. CToO performance and evaluation over in-distribution, out-of distribution (OOD) as well as cfDNA. **(a)** An overview of CToO and downstream tasks. **(b)** Macro F1 scores on in-distribution and external out-of distribution dataset. In-distribution contains 39 cell types and out-of distribution(OOD) dataset contains 6 cell types. Bars represent means; error lines indicate standard deviation. **(c)** OOD *in silico* composition before (left) and after (right) cell type enrichment. Enrichment was done with fine-tuned Pleiades 600M model over Top 1K cell type markers. **(d)** Cell type enrichment over actual cell-free DNA samples. Cell type proportion was estimated using UXM deconvolution tool. **(e)** Deconvolution benchmark against 2 well known deconvolution tools. Bars present the mean Jensen-Shannon divergence score between true cell type ratio and estimated ratios; error bars indicate the standard deviation. Root mean squared error (RMSE) is reported in Fig. S7.

We compare overall averaged F1 score of our fragment-level CToO classifier across progressively larger panels of Differentially Methylated Regions (DMRs) [77]: (i) the 25 marker regions published in the tissue methylation atlas [58], as well as the (ii) the top 100 and (iii) the top 1000 regions discovered from our training cohort (Fig. 2b). Per cell type results are detailed in Fig. S5, while the DMR calling method is described in Section 4.4. The shift in the UMAP embedding of fragment representations after fine-tuning (Fig. S4) further illustrates how the model leverages these DMRs to separate cell types in latent space.

We observe that applying stricter filtering criteria to the DMR sets enhances classification accuracy. This rigorous filtering is especially beneficial for downstream applications that require precision within narrowly defined genomic regions.

In contrast, employing larger DMR sets enables broader genomic coverage, allowing identification of a larger number of fragments originating from any cell type of interest. Across all three panel sizes, Pleiades 90M and 600*M* achieve higher macro F1 than the tuned random forest baseline (e.g. Top-25 panel: 0.67 vs 0.46; Fig. 2b). As the panel expands to Top 100 and Top 1k regions, this performance gap widens, highlighting the Pleiades’ abilities to capitalise on broader marker sets.

We then assessed performance on an unseen and independent out-of-distribution (OOD) dataset containing 6 different cell types [78] (Fig. 2b). Fig. S6 illustrates the classification F1 scores on this OOD dataset across all cell types and models. The model demonstrates comparable or improved performance across several cell types when evaluated on previously unseen data.

#### 2.4.1 Cell Type Enrichment

Having demonstrated the capability of Pleiades to accurately predict fragment-level cell type origins, we next explore the model’s potential for cell type enrichment. Enrichment involves selectively retaining fragments classified by the model as originating from a specific cell type, thereby enhancing the representation of a cell-of-interest within naturally mixed cfDNA samples. To evaluate enrichment, we utilised the OOD dataset used earlier [78] to generate an *in silico* admixture for downstream manipulation (Fig. 2c). We applied predictions from the Pleiades 600M model to select DNA fragments with the highest relative probabilities assigned to each target cell type. Due to computational constraint, we did not fine-tune Pleiades 7B.

Results revealed large increases in the target cell type proportions within the top 1K DMR regions (Fig. 2c). Notably, even rare cell types exhibited substantial enrichment: neuronal fractions increased from 7.37% to 58.5%, a 7.9-fold enhancement and hepatocytes from 4% to 73.3%, an 18.3-fold increase.

The enrichment process balanced precision, indicated by high post-enrichment cell type proportions, and recall, reflecting the retention of relevant fragments (Fig. S9). Furthermore, broader DMR regions enhance genomic coverage, allowing retrieval of more diverse fragments.

Additionally, we validated this enrichment strategy on a real-world cfDNA sample with unknown cell type composition. Employing the UXM deconvolution tool to estimate proportions before and after enrichment (Fig. 2d), we observe substantial improvements across all evaluated cell types, further confirming the effectiveness of Pleiades for cell type enrichment.

#### 2.4.2 Cell Type Deconvolution

To assess the effectiveness of our fragment-level approach, we benchmarked Pleiades against two leading deconvolution methods, UXM [58] and CelFiE [64], which infer cell type ratios based on global aggregated features from cfDNA samples.

Each experiment involved random sampling of 500,000 fragments, with known groundtruth cell type ratios and repeated 5 times with different sampling seeds. All methods require predefined marker regions, restricting the total number of fragments for evaluation. Performance is quantified using Jensen-Shannon divergence [79] and root mean squared error (RMSE). For UXM and Pleiades, we used the published Top 25 official marker set to achieve full comparability [58].

Pleiades achieved competitive overall performance compared to both UXM and CelFie (Fig. 2e, Fig. S7). Pleiades performed best in 3 of 5 categories and was tied with UXM in 1 (Fig. 2e). UXM performed best in the category of 4 cell type mixtures. CelFie did not achieve superior performance in any of the mixtures.

To assess sensitivity to rare cell types, we also performed variable-proportion spikein experiments in which individual cell types were titrated into a B cell background at frequencies ranging from 0% to 10%. We report the Pearson correlation and RMSE between predicted and true cell type proportions for the spiked-in cell types, as well as the Lin’s Concordance Correlation Coefficient (CCC) to assess calibration (concordance with the y=x line) (Fig. S8). Pleiades and UXM both outperformed CelFiE in this setting. Notably, Pleiades demonstrated strong calibration for rare cell types such as neurons and hepatocytes, as reflected by higher Lin’s CCC values (Fig. S8).

For these tasks, high-performance was achieved with the small Pleiades models. At this scale, fragment-level CToO yielded robust, generalisable cfDNA cell-type calls.

### 2.5 Applications of Pleiades to Neurodegenerative Diseases

With a deep understanding of the human epigenome and cfDNA, we hypothesised Pleiades could be applied to the detection of early-stage neurodegenerative diseases. As a modality, cfDNA promises a minimally-invasive alternative to traditional diagnostic and prognostic methods such as cerebrospinal fluid (CSF) analysis and amyloid-PET scans [80–82].

A cohort of 81 age and sex-matched patients were curated from the Cognitive Disorders Clinic at Oxford University NHS Foundation Trust, Oxford, UK (Fig. 3b). Patients were identified as having mild dementia consistent with the AD clinical syndrome, supported by an ATN-compatible biomarker profile [83]. Patients additionally underwent comprehensive cognitive evaluation and neuroimaging for completeness; full details are described in Methods, Section 4.5.1. To take advantage of Pleiades’ global genomic understanding, we utilise a whole-genome approach rather than targeted sequencing methods as previously reported [45]. cfDNA was extracted from plasma and prepared into libraries for EM-Seq using short-read sequencing at depths *>* 30x.

**Fig. 3:**
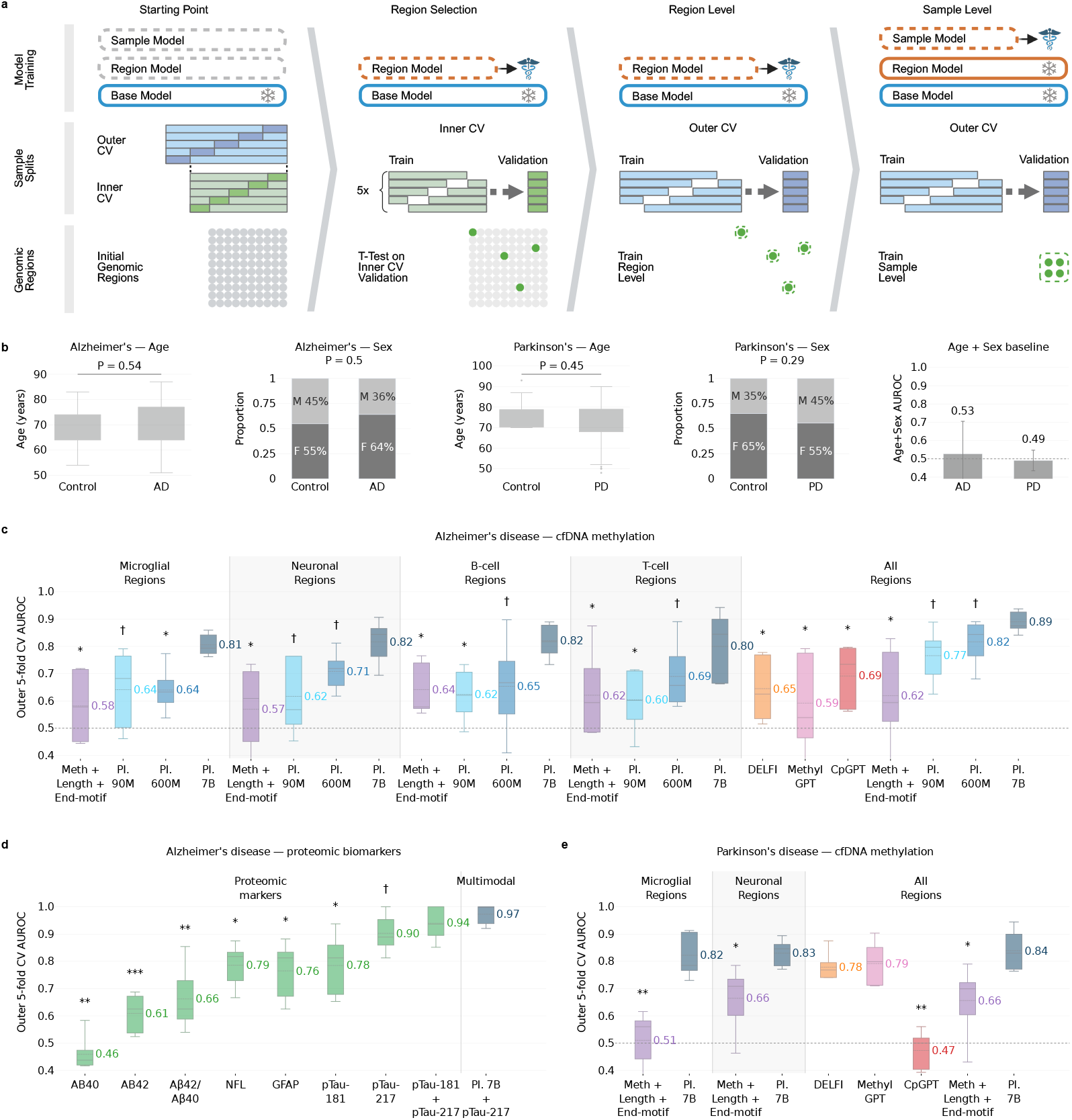
Clinical Neurodegenerative Disease Diagnosis with Pleiades. **(a)** Schematic representation of the diagnosis process using hierarchical Pleiades models. The process begins with broad genomic regions, and the entire sample set is divided into nested 5-fold cross-validation. Inner folds are used to train region-level models and perform a statistical test (T-test) to select high-performing regions on the inner validation sets. The outer-fold train sets are then used to train the region- and sample-level models, and final performance is reported on the held-out outer validation sets. **(b)** Demographic distributions of the AD and PD cohorts. Age (box plots) and sex (stacked bars; F, female; M, male) distributions are shown for the Alzheimer’s disease and Parkinson’s disease cohorts, comparing patients to age- and sex-matched controls. *p*-values were computed using a Mann-Whitney U test for age and a Fisher’s exact test for sex. The rightmost panel shows the mean test AUROC of logistic-regression baselines trained on age and sex features alone, using the same cross-validation folds as in panels (c)–(e); in both cohorts age and sex are not predictive of disease status (AD AUROC = 0.53, PD AUROC = 0.49). **(c)** Performance on clinically diagnosed AD from cfDNA. Box plots show the outer 5-fold cross-validation AUROC for all three Pleiades model sizes (90M, 600M, 7B) per cell-type marker-region set (microglia, neuron, B cell, T cell) and for the average-pooled ensemble across all regions, alongside cfDNA baseline classifiers: a combined methylation + fragment-length + end-motif linear model (per cell-type DMR set), DELFI, MethylGPT, and CpGPT. Numbers indicate mean AUROC; the dashed gray line marks chance-level performance (AUROC = 0.5). Baseline classifiers are described in Section 4.5.4. **(d)** Performance on clinically diagnosed AD from proteomics and multimodal combination. Box plots show the outer 5-fold cross-validation AUROC for the 7 individual proteomic markers (proteomic markers), the purely proteomic pTau-181 + pTau-217 combination, and the multimodal Pleiades 7B + pTau-217 model (multimodal). **(e)** Performance on clinically diagnosed PD from cfDNA, shown as in (c), for the microglia and neuron marker-region sets, the average-pooled ensemble across all regions, and the same cfDNA baseline classifiers. For panels (c)–(e), statistical significance was assessed using one-sided paired *t*-tests on the per-fold AUROCs across the five outer cross-validation folds, with Benjamini–Hochberg correction for multiple comparisons. In (c) and (e), each Pleiades model size and baseline is compared to the Pleiades 7B model within the same cell-type set; in (d), each proteomic marker is compared to the multimodal Pleiades 7B + pTau-217 model. Symbols denote adjusted significance (^*†*^ *p <* 0.1, ^∗^ *p <* 0.05, ^∗∗^ *p <* 0.01, ^∗∗∗^ *p <* 0.001); no symbol indicates a non-significant comparison.

For modelling and evaluation, clinical diagnosis was framed as an epigenomic *set* problem. Each cfDNA fragment was processed by the pretrained Pleiades base model with a trailing `[CLS]` token: the final-layer `[CLS]` embedding represents that fragment. Embeddings from the same sample are concatenated and passed to a HAT block (Fig. 1a) that builds three nested representations: (i) individual fragments, (ii) genomic regions (aggregating fragments), and (iii) the complete cfDNA sample (aggregating regions). This hierarchy mirrors biological organisation and enables region-level attribution alongside sample-level prediction.

Diagnostic performance of Pleiades for early clinical AD demonstrated high-performance (Fig. 3c). We utilised a nested cross-validation approach (Fig. 3a) as described in Methods, Section 4.5.2. Pleiades 7B achieved AUROC scores for classification of AD vs control across specific cell-type identities as follows: 0.81 for microglia, 0.82 for neuron, 0.82 for B cell, and 0.80 for T cell, on average. Smaller models, as expected, exhibited less accurate and more variable performance; Pleiades 90M achieved average AUROC scores ranging between 0.60 and 0.64 and Pleiades 600M between 0.64 and 0.71. Notably, we observed significant scaling improvements with the 7B model in both mean AUROC scores and consistency of results (Fig. 3). By combining predictions across all available cell types using average pooling, we achieved an AUROC of 0.89, suggesting the presence of complementary signals across cell types.

Further, when we trained Pleiades on experimental, non-human quality control DNA introduced within each cfDNA sample during sequencing (pUC19, Lambda DNA), all regions were rejected by our marker selection t-test with no regions deemed to contain any signal relating to clinical disease status (Fig. S10).

To investigate the biological relevance of the selected regions, we performed region-based GO enrichment analysis using GREAT [84, 85] on the pooled top-4 regions from all cell types, using the full set of starting candidate regions as background to identify processes enriched specifically among model-selected regions rather than in the candidate set as a whole (Supp. Fig. S11). The selected regions were significantly enriched for neurodevelopmental processes including nervous system development, regulation of neurogenesis, and brain development (FDR *<* 0.05; binomial test), consistent with known epigenetic dysregulation of neurodevelopmental pathways in AD [34, 86, 87].

In recent years, protein biomarkers have emerged as promising tools in the diagnosis of Alzheimer’s disease due to their strong biological relevance and established clinical significance [54, 88, 89]. In this cohort of patients, individual proteomic biomarkers achieved a range of AUROC scores: 0.46 (A*β*40), 0.61 (A*β*42), 0.66 (A*β*42/40), 0.79 (NfL), 0.76 (GFAP), 0.78 (pTau-181), and 0.90 (pTau-217) (Fig. 3d). Comparing the protein biomarkers to Pleiades, we observe improved performance (Benjamini-Hochberg adjusted *p <* 0.05) of Pleiades relative to all of the tested biomarkers individually (Table S5). Combining pTau-217 with Pleiades 7B predictions across all cell-type marker regions yielded a peak AUROC of 0.97, a +0.07 improvement over pTau-217 alone (0.90) that did not reach statistical significance after Benjamini-Hochberg correction (*p* = 0.06; *n* = 81). A purely proteomic combination of pTau-217 + pTau-181 reached an AUROC of 0.94, with no significant difference from Pleiades + pTau-217.

We further benchmarked Pleiades against four baseline classifiers applied to the same cohort and cross-validation folds: DELFI [63], a fragment-length-based linear model; a combined linear model over methylation, fragment-length, and 5^*′*^ end-motif features (Meth + Length + End-motif), in the spirit of [90]; MethylGPT [91], and CpGPT [16]. These baselines span both major cfDNA feature classes —fragmentomics and methylation — and range from established linear approaches to recent foundation models, providing a broad comparison. All baseline methods underperformed Pleiades 7B (Fig. 3c), consistent with the hypothesis that Pleiades’ fragment-level representations capture disease-relevant epigenomic signals beyond what CpG-level methylation summaries or fragment length distributions alone can provide.

To assess the transferable capabilities of Pleiades across conditions, we apply the model to the early detection of PD. 160 human plasma samples from patients with PD and age- and sex-matched controls were procured from a commercial supplier (Fig. 3b). Samples were processed in-house identically to AD samples. Microglial region AUROC scores for Pleiades 7B models are on average 0.82, and neuronal 0.83 across 5 outer folds (Fig. 3e). The average pooled version of these models achieves AUROC of 0.84. The same baselines evaluated for AD were applied to this cohort; all underperformed Pleiades 7B (Fig. 3e)

All disease diagnosis experiments used a nested 5-fold cross-validation, with outer folds giving error bars (Fig. 3c-e). A detailed example of the methodology given an outer fold is described in Supp. Fig. S10.

Overall, multi-cell-type cfDNA ensembles using Pleiades matched or exceeded the seven proteomic markers tested. Multi-modal combination with pTau-217 reached a peak AUROC of 0.97.

### 2.6 Mechanistic Interpretability of Pleiades’ Alzheimer’s Disease Prediction

We next sought to identify the biological signals driving Pleiades’ Alzheimer’s disease predictions using mechanistic interpretability. Interpretability is particularly valuable for clinical applications, where understanding model behaviour may be of high importance to clinicians and regulators.

Applying a top-*K* sparse autoencoder (SAE) [92] to the Pleiades 7B `[CLS]` embeddings yielded a sparse dictionary of 761 alive features (Methods, Section 4.6.2). Ranking these features by gradient attribution to the AD logit [93] showed that AD prediction concentrated in a compact subset rather than drawing on the full dictionary: the top 20 of the 761 features accounted for 78.4% of the total gradient attribution (Fig. 4a, top). Single-feature zero-ablation confirmed this independently; the top-ranked features produced the largest shifts in the AD logit (mean |Δlogit| = 0.26 across the top 20), and the two measures largely agreed on which features drove prediction (Fig. 4a, bottom; extended across the top-100 alive features in Supp. Fig. S12b; Methods, Section 4.6.3). Given this concentration, we next asked what biological information these top-20 features encode.

**Fig. 4:**
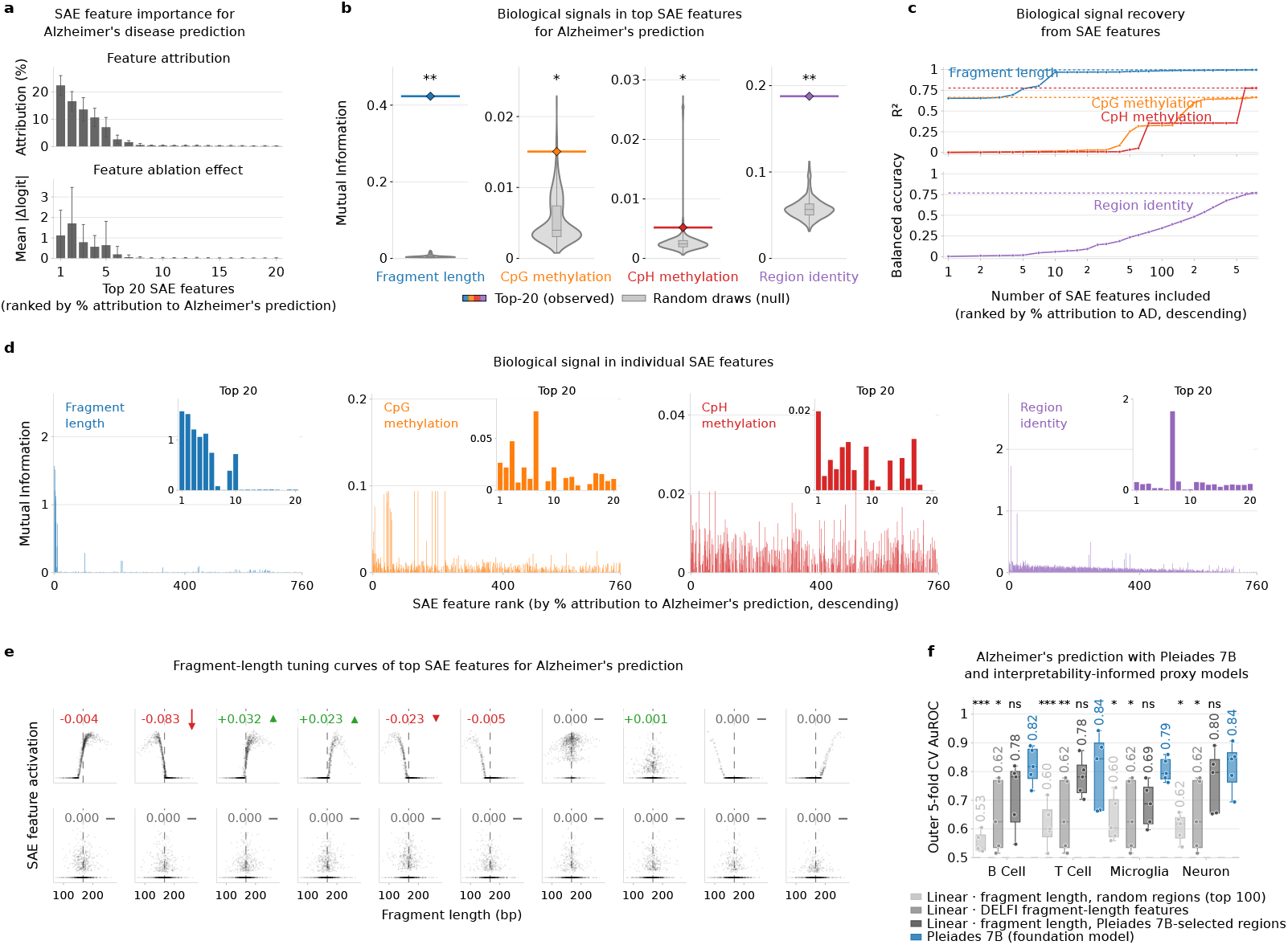
Mechanistic Interpretability of Pleiades AD Classification. **(a)** Top 20 sparse autoencoder (SAE) features driving Alzheimer’s disease prediction. SAE features are ranked by gradient attribution to the Alzheimer’s prediction logit; x-axis shows the top-20 attributed features (sorted high to low). This order is also used in subsequent panels. The top y-axis shows the % gradient attribution for each SAE feature. The bottom y-axis shows the mean per-individual |Δlogit| under single-SAE-feature zero-ablation. Error bars across cell type *×* fold. **(b)** Mean mutual information (bits) for the top-20 SAE features by AD attribution (diamond/line) versus a null from 1,000 random 20-feature draws (grey violins), for fragment length, CpG/CpH methylation, and region identity. Stars denote BH-adjusted permutation *q*-values: ^∗∗∗^, *q <* 0.001; ^∗∗^, *q <* 0.01; ^∗^, *q <* 0.05. Independent *y*-axes. **(c)** Number of SAE features needed to reconstruct biological signals with linear regression (or logistic regression for classification tasks). held-out score (*R*^2^ or balanced accuracy) is plotted against SAE feature-set size, with features added by descending attribution score. Dotted lines indicate asymptotes for each feature. **(d)** Mutual information between individual SAE features (sorted by descending attribution score) and biological signals: fragment length, region identity, and methylation ratio (CpG, CpH). Insets show mutual information for the top-20 SAE features. **(e)** Activations of the top-20 SAE features as a function of fragment length. The effect of ablating each feature on Alzheimer’s prediction is shown in the arrows annotating each plot. Dashed vertical lines indicate the mean fragment length across the dataset. **(f)** AD-classification AUROC across four cell types (mean *±*1 SD over five folds) for Pleiades 7B and three linear models based on fragment length features. ‘Fragment length, random regions (top 100)” uses median fragment length in randomly sampled regions from the top 100 ones of that cell type, “DELFI fragment-length features” uses fragment length features in 5MB windows following the DELFI method [63], and “fragment length Pleiades 7B-selected regions” uses median fragment length in regions selected from the cross-validated folds in Figure 3. Finally, we show the “Pleiades 7B” model as in Figure 3. Cross-model significance was assessed per cell type by a one-way repeated-measures ANOVA (cross-validation fold as the repeated factor) followed by Dunnett post-hoc tests against Pleiades 7B as the control (Supp. Table S6).

We hypothesised four candidate types of biological signal that Pleiades may have learned about cfDNA fragments: length, CpG methylation ratio, CpH methylation ratio, and genomic region identity. For each signal, we compared the mean mutual information (MI) of the top 20 SAE features ranked by attribution to Alzheimer’s disease (AD) prediction against a null distribution from 1,000 random draws of 20 SAE features (alive features outside the top 20). The top-20 feature set carried significantly more signal than randomly drawn 20-feature sets across all four signals (one-sided empirical permutation test, Benjamini–Hochberg corrected over the four signals). Mean mutual information for the top-20 set versus the random null was 0.42 vs. 0.005 bits for fragment length (*q* = 2.0 × 10^−3^), 0.015 vs. 0.005 bits for CpG methylation ratio (*q* = 2.0 × 10^−2^), 0.0052 vs. 0.0030 bits for CpH methylation ratio (*q* = 4.0 × 10^−2^), and 0.19 vs. 0.058 bits for genomic region identity (*q* = 2.0 × 10^−3^) (Fig. 4b).

We further explored whether this difference could be explained by how each type of biological signal is encoded in the SAE features, reasoning that some signals may be concentrated in a small number of SAE features while others may be more diffusely spread across many features. We found that fragment length could be predicted well from a small number of SAE features; the top-20 attribution-ranked features alone recovered 96.8% of fragment-length variance (*R*^2^ = 0.968, approaching the all-feature ceiling of 0.996) (Fig. 4c). Regarding region identity, the top-20 features reached 9.4% balanced accuracy across 354 classes (33× the 0.28% chance rate), rising to 76.9% only when all 761 active features were included. In the case of methylation, the top-20 features recover up to 3% of methylation-ratio variance (CpG *R*^2^ = 0.029; CpH *R*^2^ = 0.008). Predicting CpG methylation ratio from SAE features required ≈500 features to saturate and reach 95% of its 0.66 asymptote (Fig. 4c), indicating that the methylation signal is present but spread diffusely across the SAE features.

We next zoomed into the individual SAE features, to identify which specific features captured which biological signals. Features at ranks 1-6, 9 and 10 captured fragment length and feature at rank 7 captured region identity (Fig. 4d). Methylation signal, particularly CpG, was spread across many of the top features at low per-feature mutual information (*MI <* 0.1 for CpG and *MI <* 0.021 for CpH) (Fig. 4d), with the rest of the features carrying information diffusely, which is consistent with both of the previous results: the top-20 set carrying more methylation signal than the random null (Fig. 4b) and with the many features required to reconstruct it (Fig. 4c).

Visualizing the activations of the top-20 SAE features as a function of fragment length (Fig. 4e), we observed that fragment length features are clearly grouped into two categories of short and long fragments (e.g. the features at ranks 1, 3, 4 and 10 capture long fragments, whereas those at ranks 2, 5, 6 and 9 capture short fragments). Consistent with this, fragment length organised the leading principal components of the model’s activations into a smooth trajectory (Supp. Fig. S12a), marking it as a dominant axis of Pleiades’ representation and alone accounted for 49.66% of final-layer `[CLS]`-embedding variance.

To test whether the top-20 SAE features causally drive Alzheimer’s prediction, we zero-ablated each feature and measured the resulting change in predicted *P* (AD). On average across features and regions, ablating short fragment tuned features lowered *P* (AD) by a mean of 0.028, whereas ablating long fragment tuned features raised it by a mean of 0.013 (arrows in Fig. 4e). The two largest effects came from the rank-2 and rank-3 features: ablating the rank-2 feature (activation-weighted mean ≈147 bp) lowered *P* (AD) by 0.083, while ablating the rank-3 feature (≈ 183 bp) raised it by 0.033 (pooled across regions). Amplifying each feature to 30× its baseline activation produced the opposite effect: shortfragment-length features raised *P* (AD) and long-fragment-length features lowered it, with the polarity reversing for some cell-type-specific regions (Supp. Fig. S13). Steering and ablation moved *P* (AD) in opposite directions across the 20 features (Supp. Fig. S12c).

Finally, we tested our mechanistic interpretability findings by asking whether a simple, interpretable proxy model can recapitulate Pleiades’ Alzheimer’s prediction performance. We compared Pleiades 7B against three logistic regression baselines (Fig. 4f). The first two used only the median fragment length per region, and differ only in the regions provided to the model: either four regions drawn at random from the dataset, denoted “random regions”, or the top-4 attribution-selected regions in each fold, denoted “Pleiades regions”. The third one uses DELFI fragment-length features, denoted “DELFI features” [63]. This design let us isolate the contribution of genomic identity to the performance of these simplified models.

The Pleiades-regions linear baseline outperformed both the DELFI and the random regions ones, achieving a mean AUROC across cell types of 0.74 compared with 0.65 and 0.60, respectively. By cell type, a repeated-measures ANOVA found the four models differed significantly in every case (all *p <* 0.025); Dunnett tests against Pleiades 7B placed both the DELFI and random-regions baselines significantly below it (*p* ≤ 0.043 and *p* ≤ 0.018). Across the four cell types, the Pleiades-regions baseline and Pleiades 7B achieved AUROCs of 0.72 vs. 0.82 for B cells, 0.78 vs. 0.80 for T cells, 0.68 vs. 0.81 for microglia, and 0.76 vs. 0.82 for neurons (Dunnett test against Pleiades 7B: *p* = 0.19, 0.89, 0.096, and 0.74, respectively; Supp. Table S6).

## 3 Discussion

Our work demonstrates that modelling both DNA and methylation unlocks capabilities beyond DNA-only language models. Methylation represents a critical feature set of the epigenome, the dynamic set of modifications of DNA that extensively influence cellular identity, function, and change throughout age and disease [26, 27, 29]. The Pleiades series of models were created to capture these changes and showcase the utility epigenetic pretraining across technical, biological, and clinical applications.

Pleiades was trained upon a unique corpus of human DNA and methylation sequences totalling 1.9T tokens. This corpus includes a comprehensive atlas of the human methylome, spanning 39 cell-type groups [58]. Notably, the diagnostic fine-tuning presented in this work utilises EM-Seq data, demonstrating that representations learned from a predominantly WGBS pretraining corpus can generalize to another methylation sequencing technology.

Pleiades spans three parameter sizes - 90M, 600M and 7B parameters - and utilises alignment embeddings to build a latent understanding of the human genomic space. We assessed Pleiades using the popular Nucleotide Transformer benchmark, which we modified to remove a bias that may influence evaluation. We showed that all Pleiades model sizes exhibit state-of-the-art macro performance on this benchmark. Small Pleiades models outperform DNA-only models with many-fold higher in parameter count, and Pleiades 7B achieves MCC of 0.98. It is likely that incorporation of high-quality methylation data during pretraining will improve the performance and capabilities of genomic language models in many downstream evaluations, perhaps beyond those presented in this work. Uniquely, Pleiades 7B additionally demonstrated few-shot learning capabilities on this benchmark. Beyond fine-tuning, frozen Pleiades 7B embeddings outperformed recent genomic foundation models on histone-mark prediction across the Nucleotide Transformer and ENCODE benchmarks, under both linear-probe and 1-shot evaluation. Under the linear probe, Pleiades 7B surpassed AlphaGenome on the ENCODE histone-modification tracks (median MCC 0.88 versus 0.64) even when AlphaGenome was given an 8 kb context, eight times the 1 kb window Pleiades uses, and despite AlphaGenome being pretrained to predict ENCODE histone-modification tracks. Pleiades learns only from DNA sequence and its methylation, never from these tracks. That it nonetheless outperforms a model trained directly on them highlights the value of methylation-aware pretraining.

In our work, we are interested in the biology of the brain in age and disease. Historically, this has been prohibited by the lack of ground-truth data and the presence of complex clinical phenotypes [40–42]. In oncology and other complex diseases, applications of epigenetics have enabled the discovery of both novel diagnostics and therapeutic targets [57, 59, 60]. With recent research identifying a key role for epigenetic disruption early in neurological pathology [32], we hypothesised a role for epigenetics to dissect the complexity of neurological disease.

We therefore explored a role for Pleiades as a downstream discovery tool for neurodegenerative conditions. We first showcased a variety of applications for the model series on cfDNA, starting with their generative capabilities. Pleiades 7B generated *in silico* fragments of cfDNA with high accuracy across methylation context and fragment size. This indicates the largest model in the series has developed a good latent understanding of the statistical properties of human methylation and cfDNA. Downstream evaluation of *in silico* generated fragments in clinical tasks will be necessary to confirm utility, which, if demonstrated, may bolster small clinical datasets, calibrate diagnostic assays, and/or create controlled settings for studying cell-specific epigenetic changes.

Pleiades was additionally able to identify the cellular origin of plasma-derived cfDNA, matching state-of-the-art deconvolution methods. Crucially, Pleiades operates at the individual fragment level rather than producing only aggregate sample-level estimates: while UXM and CelFiE estimate what fraction of cfDNA comes from a given cell type, Pleiades can identify *which* individual fragments originate from that cell type. This per-fragment resolution enabled selective enrichment of rare but informative fragments, as demonstrated by the 7.9-fold enrichment of neuronal fragments and 18.3-fold enrichment of hepatocyte fragments in our *in silico* experiments. In a clinical context, such enrichment could enhance the sensitivity of downstream diagnostic or monitoring assays by concentrating disease-relevant signal from brain-derived cfDNA, which is present at very low fractions in plasma. Future work will characterise the clinical utility of cell-type-based enrichment in prospective neurodegenerative disease cohorts.

We then demonstrated Pleiades’ ability in biomarker discovery, given a starting set of genomic regions. Our results support the high-performance of both cfDNA and proteomic approaches independently. We suspect a combination of foundation modelling for both to be advantageous, for improved AUROCs and further stratification or subtyping of disease. It is possible that the underlying mechanisms for both proteomic and epigenetic signatures identified are via different mechanisms, e.g. neuronal loss and microglia-induced inflammation for cfDNA and protein oligomerisation/aggregation for proteomic markers. Downstream work would benefit from interrogation of the methylation subtypes and genomic regions that drive model decisions. We took a first step in this direction by mechanistically dissecting Pleiades’ AD predictions, grounding the model’s latent features in interpretable biological signals rather than treating the classifier as a black box. Across complementary angles, such as attribution, causal perturbation, and surrogate modelling, these analyses converged on a consistent account, relating the predictions to cfDNA fragmentomic properties at genomic regions the model had itself selected. The same fragment-length classifier reached an AUROC of 0.74 over the Pleiades-selected regions but only 0.60 over randomly chosen ones, showing that the model pinpoints where the diagnostic signal resides. The fragment-length surrogate restricted to these model-selected regions recovers most but not all of Pleiades’ performance (mean AUROC 0.74 vs. 0.81), with the residual likely reflecting nonlinear or more diffusely encoded signal such as methylation. These analyses are computational, however, and establishing any such property as a usable biomarker will ultimately require experimental verification in the laboratory, an essential direction for future work.

### Limitations

Despite these advances, several constraints remain:

1. **Limited training data**. Pretraining relied solely upon methylated DNA from healthy human tissues and cfDNA. The use of predominantly healthy tissue samples for pretraining is by design: the objective is to capture the general distribution of epigenetic patterns across normal cell types, providing broad, unbiased foundational representations onto which disease-specific signals are learned during task-specific fine-tuning. The absence of other epigenetic marks and non-human genomes curtails multi-omic and cross-species generalisation. As shown in Section 2.5, multimodal combination of cfDNA with the plasma protein marker pTau-217 yields a higher peak AUROC than either modality alone, suggesting that broader multi-omic data at both pretraining and fine-tuning stages could yield further gains. Recently published work indicated strong potential for the inclusion of large, multi-parametric proteomic information for downstream diagnostics and prognostics for neurological conditions [44, 94, 95]. In addition, our work has focussed on whole-genome sequencing of cfDNA (30-50x). While this allows for a global view of a patient biosample, it does increase the risk of over-fitting. Additionally, drop in whole-genome sequencing costs does require consideration of much higher depths of sampling (i.e. *>*1,000x). Target approaches such as bespoke methylation panels may also be beneficial.
2. **Cohort size and diversity**. The clinical cohorts assessed for downstream diagnosis are modest (81 AD, 160 PD) and were recruited primarily from individual centres within the United Kingdom. While the AD cohort was prospectively recruited and characterized with ATN biomarker profiling, and the PD cohort was independently procured with clinical diagnostic confirmation, neither cohort captures the demographic diversity necessary to establish clinical generalisability. Population stratification in DNA methylation patterns has been documented across ancestries [96], and differences in cfDNA fragment profiles may similarly vary with genetic background, comorbidities, and environmental exposures. Prospective validation in larger, multicentre cohorts spanning diverse ancestries, geographic regions, and healthcare settings will be essential. Furthermore, larger additional cohorts would also increase statistical power to confirm whether multi-modal cfDNA + pTau-217 integration significantly outperforms proteomic combinations alone. Until such validation is complete, the diagnostic performance reported here should be interpreted as a proof-of-concept demonstrating the potential of epigenomic foundation models for neurodegenerative disease detection, rather than as definitive clinical performance estimates.
3. **Further interpretability and clinical actionability**. We identified multiple marker regions within the genome and specific cfDNA properties that drive the diagnostic predictions. However, the model does not yet fully annotate *what* biological mechanisms underlie the identified epigenetic signals, nor *why* specific regions are selected across cell types. Sparse autoencoders yield useful but approximate and non-identifiable decompositions of model representations and are not guaranteed to be robust [97]. Feature attribution methods also offer approximate evidence of feature influence rather than exact causal accounts [98]. Moreover, the candidate signals we tested are neither exhaustive nor fully independent, and the fragment-length proxy recovers some but not all of Pleiades’ performance, leaving additional, likely nonlinear, signal uncharacterised. For clinical translation, model outputs would need to be integrated alongside established diagnostic criteria (e.g., ATN status for AD, Movement Disorder Society criteria for PD), proteomic biomarkers, neuroimaging, and cognitive assessment [80, 83]. Calibrated prediction probabilities with explicit confidence intervals, and prospective validation of clinical decision thresholds, would be required before deployment, as well as clear generalisability. We envision Pleiades-based cfDNA analysis as a component of a multi-modal diagnostic pathway, contributing complementary epigenomic information rather than serving as a standalone decision tool.

### Future Work

Multi-modal information will improve the capabilities of Pleiades for both biomarker discovery and future mechanism identification for downstream targets. Expanding Pleiades to additional epigenomic modalities (ATAC-seq, ChIP-seq, etc.) alongside the integration of transcriptomic and proteomic data may improve diagnostic and prognostic capabilities. Incorporation of high-quality single-cell brain atlases of ageing and degeneration could yield a multi-omic brain foundation model [76]. Incorporating set-level pretraining objectives alongside the autoregressive loss, together with architectural advances for representing very large sets, may produce more expressive sample-level embeddings and boost diagnostic and generative performance. Finally, different strategies for incorporating alignment information, such as using CIGAR strings instead of per-base embeddings, warrant further exploration.

Building upon results observed when scaling to 7B parameters, we also plan to investigate larger foundation models and memory-efficient transformer variants that accommodate substantially longer context windows — potentially on the order of megabases — so that Pleiades can tackle genome-scale set problems end-to-end. Finally, we seek to develop a dedicated interpretability pipeline aiming to turn model attributions into mechanistic insights and clinically actionable biomarkers. In a clinical setting, such attributions — combined with calibrated uncertainty estimates — could support clinician decision-making by identifying which genomic regions and fragment-level features contribute most to a given patient’s risk score, enabling targeted follow-up investigations.

Multi-modal foundation modelling for biology offers promise to enable precision medicine and unlock novel insights for complex diseases. Pleiades establishes the first step and displays the effectiveness of jointly modelling DNA and methylation in a unified, general-purpose foundation model. Our work lays the groundwork for accelerated biomarker discovery, and deeper, interpretable mechanistic insights into the genomic regulation of brain ageing and disease. Near-term, effective staging and subtyping of disease would improve clinical outcomes and could enable a molecular classification of neurological disease. Longer-term, wider expansions of modality and improvements in training architecture and methodology would encourage the creation of a brain foundation model for discovery of novel mechanisms and targets for precision therapies.

## 4 Methods

### 4.1 Pretraining

#### 4.1.1 Architecture and Pretraining Procedure

The Pleiades base model is an auto-regressive language model based on the generative pretrained transformer architecture [99]. It uses character-level tokenisation that represents each nucleotide with a capital letter (`A, C, T, G`) and methylation as (`<m>`). For the activation function of the Multi Layer Perceptron (MLP) layers, we used squared ReLU [100] for three primary reasons: (a) it is comparatively performant on several language-related tasks [101, 102]; (b) it can produce sparse representations; and (c) it has been empirically shown to be the most efficient activation function for sparse LLMs [103]. Rotary positional embeddings (RoPE) [104] were utilised to represent the relative position of input tokens.

The models are optimised using the AdamW algorithm [105] with a learning rate of 10^−4^ and a cosine annealing learning schedule. In order to speed up training and reduce memory usage, we used bfloat16 mixed precision [106]. Full hyper-parameters are specified in Table 1.

**Table 1:**
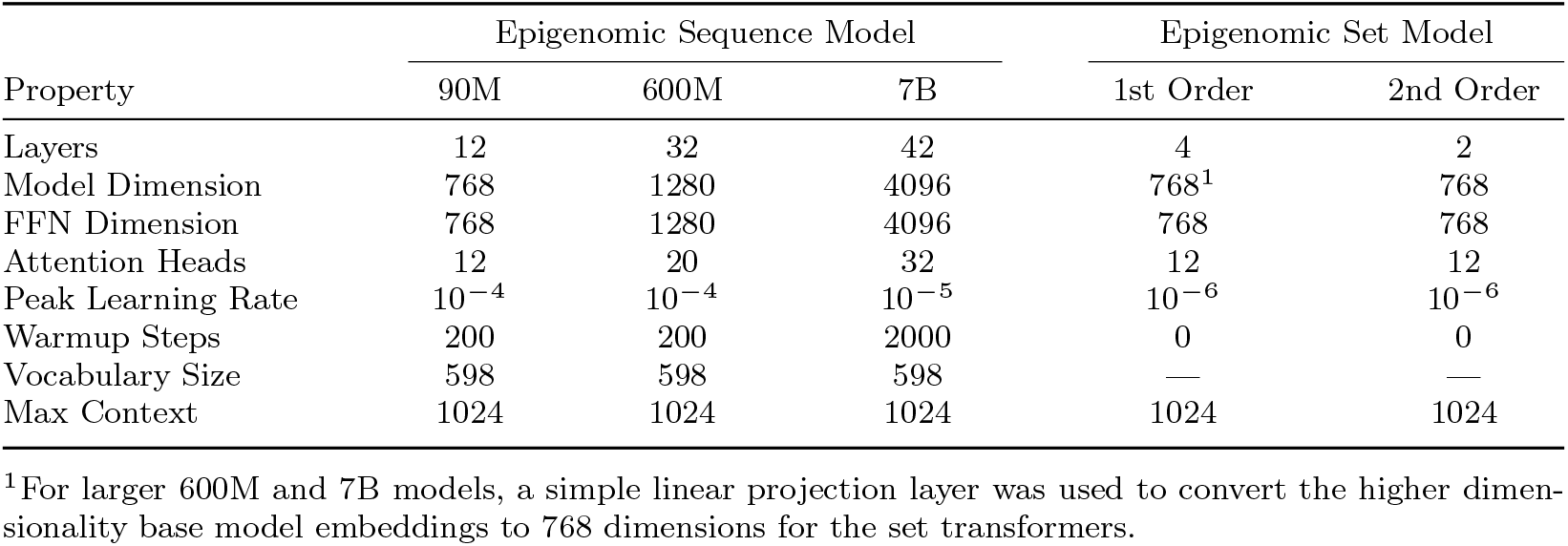
Pleiades Architecture Details.

For pretraining, we used the standard cross-entropy objective function. Given a sequence *x* = (*x*_1_, … , *x*_*T*_), the autoregressive language model is trained to minimise the negative log-likelihood:

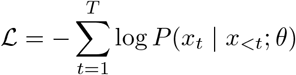

where *P* (*x*_*t*_ | *x*_*<t*_; *θ*) is the model’s predicted probability of token *x*_*t*_ given the previous context *x*_*<t*_.

The input to the model during pretraining consisted of sequences from various sources including the human genome, specific cells, and plasma. The variability of the type of sequences was introduced to the model with special tokens at the beginning and the end with `<dna>,<mdna>` and `<cfdna>` for pure nucleotide, sequences that include methylation, and cfDNA fragments respectively. If the cell type of origin is known, a `<cell type>` token is used followed by the name of the cell type and the closing token `</cell type>`. Table 2 shows how these three different modalities would look like as sequences of tokens.

**Table 2:**
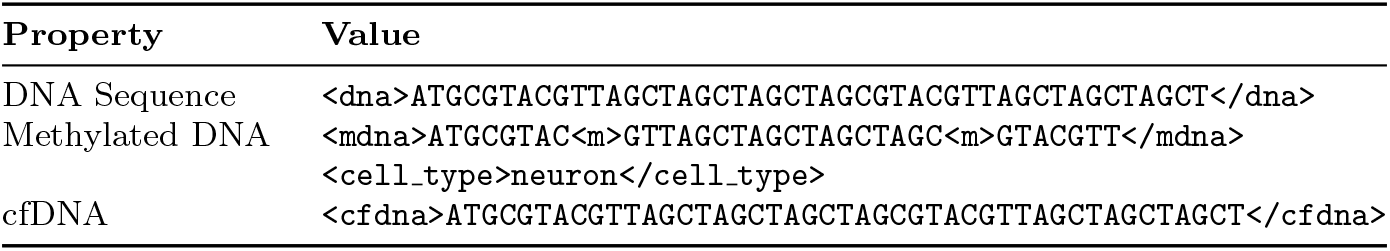
Examples of DNA Representations.

#### 4.1.2 Alignment Embeddings

Alignment Embeddings (AEs) encode genomic location by explicitly defining the chromo-some number and precise position for each nucleotide, computed with reference to the Concise Idiosyncratic Gapped Alignment Report (CIGAR) strings against the GRCh38 reference genome. Given that the human genome comprises over 3 billion base pairs, positional information can span extensive ranges, exemplified by the largest contiguous segment (chr1) containing approximately 249 million base pairs [107]. To be equally sensitive to changes in genomic position in the largest to the smallest scale, we segmented the position within the chromosome into three distinct parts: millions, thousands and ones. Consequently, each nucleotide was represented by a set of four tensors corresponding to the chromosome and segmented positional values:

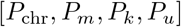

with constraints

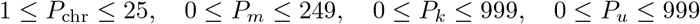

Note that we encode chromosome X, Y, and the mitochondrial chromosome as 23, 24 and 25 respectively. These components are individually embedded via learned embeddings as follows:

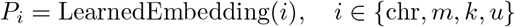

#### 4.1.3 Set-level modelling with a multi-tier Hierarchical Attention Transformer

Many clinical and biological applications rely on *sample-level* information drawn from sets that contain 10^8^ − 10^9^ cfDNA fragments. To obtain a compact representation of such large sets we place a **Hierarchical Attention Transformer (HAT)** [71] on top of the frozen sequence-level Pleiades model (Fig. 1a).

##### Architecture

Each fragment is first processed by the base decoder; the corresponding contextual `[CLS]` embedding forms the input to the HAT. A single HAT block attends over this collection of embeddings and emits a *set token* that summarises the group. We *stack N* such blocks, so that the output token of tier *k* serves as one element in the input set of tier *k*+1. This recursive design yields successively higher-order representations — fragment → region → sample — while keeping memory requirements fixed.

##### Data flow

1. **Tier 0**: sequence decoder produces one `[CLS]` vector per fragment.
2. **Tier 1**: a HAT encoder attends over all fragment vectors within a genomic window and outputs a region vector.
3. **Tier** 2 … *N* : region vectors are concatenated and passed through additional HAT encoders; the final `[CLS]` token forms the input to the task-specific head.

##### Advantages

The multi-tier scheme (a) preserves information at multiple genomic scales, (b) accommodates arbitrary set sizes, and (c) distinguishes fragments from different loci without enlarging the transformer context window. All hyper-parameters are listed in Table 1.

#### 4.1.4 Pretraining Datasets

The pretraining data corpus is a compilation of four different sources: (1) the DNA methylation atlas of normal human cell types [58]; (2) WGBS of cfDNA from healthy individuals [64]; (3) EM-Seq of cfDNA from healthy individuals; (4) and a genomic graph of the 1000G dataset [65], a human WGS dataset that captures high haplotypic variation. The corpus includes both single-end and paired-end sequencing reads. For single-end reads and haplotype reference reads, each read is treated as an individual fragment. For paired-end reads, we merged overlapping read pairs into a single fragment to increase the effective sequence context and reduce redundancy.

The DNA methylation atlas is a comprehensive WGBS dataset of 39 human cell types from 205 healthy tissue samples. It provides high-resolution methylation maps at the fragment level rather than just individual CpG sites. Loyfer et al. (2023) [58] identify over 1000 cell type specific Differentially Methylated Regions (DMRs), focusing on uniquely unmethylated/methylated regions, which usually reside in enhancers and contain binding sites for tissue-specific transcriptional regulators. These DMRs are consistent between individuals, reflecting the cell lineage and cell type specific programmes. To leverage this dataset for pretraining, we prioritise CpG-rich regions by including all CpG islands, shores, and shelves, while downsampling open sea regions to 5%.

We utilise cfDNA datasets prepared with experimental methods to capture methylation information. Specifically, we include 10 healthy samples from Caggiano et al. (2021)’s [64] study generated via WGBS alongside 10 healthy samples from our clinical samples generated via EM-Seq. In both cases, the data format is FASTQ and is processed using a modified MethylSeq pipeline [108] with Bismark [109] as the aligner. After we generated the downstream BAM files, we followed the same pre-processing and filtering steps as the DNA methylation atlas [58].

Following a similar process to that in Dalla-Torre et al. (2024) [15], we combined the human reference genome (GRCh38) with phased haplotype variant data from the 1000 Genomes Project (release version 20220422) to construct chromosome-specific variation graphs [65]. The construction process used VG (version 1.62.0) [110], employing the “construct” command to integrate sequence and variation data. Each variation graph was indexed using the 1000G preset, ensuring the retention of haplotype-specific pathways. To analyse localised genomic variation, a sliding window approach was applied to each variation graph. A window size of 1 million base pairs was defined and the window moved incrementally along reference positions. For each window, 40 haplotype paths were randomly sampled from the preserved haplotype pathways. This subsampling was conducted to represent diverse genomic contexts within a manageable computational framework. Simulated sequencing reads were generated from the subsampled haplotype paths for each window. Read simulation was performed to achieve approximately 20x coverage per haplotype. Each read was assigned a random start position within the window and read lengths were restricted to a range of 500 to 1000 nucleotides. These parameters were selected to mimic the characteristics of real-world sequencing while maintaining high coverage for downstream analysis.

#### 4.1.5 Computational Resources

We pretrained Pleiades 7B for approximately 10 days on 256 H100 GPUs (32 nodes × 8). For the Unbiased Nucleotide Transformer benchmark, we fine-tuned between ∼4 minutes and ∼1 hour (depending on the task) on a single H200 GPU. Pleiades 600M was trained for ∼18h on 8 H200 GPUs (1 node) for Top 25 regions, ∼11h on 64 H200 GPUs (8 nodes) for Top 100 regions and ∼22h on 64 H200 GPUs (8 nodes) for Top 1000 regions. Finally, for our disease diagnosis task we performed ∼17h of reconstruction-loss training of Pleiades 7B on 128 H100 GPUs (16 nodes) followed by disease diagnosis fine-tuning for ∼8h per cell-type region on 32 H200 GPUs (4 nodes).

### 4.2 Nucleotide Transformer Benchmarks

In order to test the performance of Pleiades and the baseline models on the Nucleotide Transformer benchmarks [15], we first had to create a newly randomised version of the Nucleotide Transformer benchmarks to counter the effects of the strong bias in positions of negative sequences in the dataset. Supplementary Section S1 and Fig. S2 explain the problem within the official Nucleotide Transformer benchmarks and our proposed solution in detail.

The next step was to fine-tune all models for exactly five epochs on the revised Unbiased Nucleotide Transformer Benchmark train set and measure the MCC. We note that although Pleiades is able to model both pure nucleotide sequences, as well as sequences that include methylation, for these benchmarks we fine-tune on input DNA sequences (without any methylation information). The reported values are the test set MCC at the end of training.

For Pleiades, a `[CLS]` token was appended to the end of the input sequence and its embeddings were directly fed to a classification head. This classification head is a simple 2−layer MLP with a ReLU non-linearity in between. The first layer has the same input and output dimension which equals the base model’s hidden dimension size. The second layer projects down to as many dimensions as the task has labels, 2 for binary classification tasks and 3 for the tasks that have 3 labels, *Splicing (All)* and *Enhancers (Types)*. Table 3 shows the exact hyperparameters used for fine-tuning Pleiades models on NT tasks. All Pleiades models were fine-tuned on H200 GPUs.

**Table 3:** Pleiades Hyperparameters for Unbiased Nucleotide Transformer Benchmarks.

| Property | 90M | 600M | 7B | 7B High LR<sup>1</sup> |
| --- | --- | --- | --- | --- |
| Peak Learning Rate | $10^{-4}$ | $10^{-4}$ | $10^{-6}$ | $6 \times 10^{-5}$ |
| Global Batch Size | 48 | 288 | 288 | 96 |
| Unfrozen Layers | *All* | *All* | *All* | Last 2 Layers |

<sup>1</sup>This setting was only used for the two tasks *Splicing (All)* and *Promoter (TATA)*.

Pleiades 90M and 600M were fine-tuned for an epoch on the DNA-only portion of our pretraining dataset, to bring their representations closer to pure DNA before fine-tuning for the benchmark tasks. This was not performed for Pleiades 7B.

Both baseline models were fine-tuned using their published code and with default hyper-parameters. DNABERT-2 was fine-tuned using the fine-tune script in this official github page with learning rate 10^−4^ on V100 GPUs with the maximum batch size that would fit per device. NT 2.5B MS was fine-tuned using LORA and with a learning rate 5 × 10^−4^ on H200 GPUs.

#### 4.2.1 Frozen-embedding evaluation

We evaluated frozen embeddings from Pleiades and the baseline models on three datasets, with no weight updates to any model. The test split was chromosomes 20 and 21. The train split was all remaining autosomes plus chromosome X. Chromosome Y was excluded. Each window was stored as a DNA sequence with a class label.

The ENCODE histone dataset comprised 884 histone ChIP-seq tracks. Each positive window was a 1,000 bp interval centred on the peak anchor of an ENCODE narrowPeak call, taking the 500 bp on either side of that anchor. Negative windows were not peak-anchored: they were random 1,000 bp genomic windows that did not overlap any positive peak across the union of all tracks, did not overlap the ENCODE hg38 blacklist, had RepeatMasker coverage at most 50%, and contained no ambiguous bases. Up to 300 positive and 300 negative windows were sampled per track per split. Sequences were taken from the hg38 reference. The NT histone (10 tasks) and NT regulatory (8 tasks) datasets were drawn from the Unbiased Nucleotide Transformer Benchmark, with balanced per-class subsampling of approximately 500 examples per class and the same chromosome 20/21 test split.

Each window sequence was passed once through each un-finetuned model in evaluation mode under `torch.no grad`, and a single mean-pooled vector was stored. For Pleiades 7B, the embedding was the mean of the final hidden states over the real, non-special, non-padding tokens (dimension 4,096). For NT MS 2.5B, it was the mean over the last hidden layer across the 6-mer tokens. For Evo 2, hidden states were taken at a fixed intermediate layer and averaged over the sequence tokens. For AlphaGenome, the one-hot-encoded sequence was passed through the trunk at 128 bp resolution and the embedding was the mean over the bins overlapping the window. The CpG/GC baseline was a two-dimensional vector per window: the CpG ratio (count of `CG` divided by length minus one) and the GC fraction.

Pleiades operated on a 1 kb context window throughout. Evo 2 and AlphaGenome were additionally evaluated at an 8 kb context, where the window was expanded symmetrically around its midpoint to 8,000 bp of hg38 sequence, embedded in full, and averaged only over the tokens of the original central window. AlphaGenome was evaluated at 2 kb in place of 1 kb. Only Evo 2 and AlphaGenome were evaluated beyond 1 kb.

Embeddings were scored under two regimes. For both, a standard scaler was fit on the full label-free training pool and applied to the train and test embeddings. In the 1-shot regime, one labelled example per class was drawn at random as a class prototype and each test window was assigned to the nearest prototype by squared Euclidean distance in standardised space; this was repeated for five random draws and the mean MCC reported. In the linear-probe regime, an L2-regularised multinomial logistic regression (*C* = 1.0, balanced class weights) was trained on the full training set. Across models, the labelled and test windows were restricted to coordinates present in every model’s embedding table and drawn once per task and repeat, so that all models were scored on the same windows. A task was scored when its shared test set contained at least 50 windows for ENCODE or 20 windows for the NT tasks, with at least two classes present in both splits. MCC was computed with `scikit-learn`.

For each dataset and regime, Pleiades 7B was compared against each baseline by pairing the per-task or per-track MCC values and applying a two-sided paired *t*-test, with Benjamini–Hochberg FDR correction within each dataset and regime.

### 4.3 Epigenomic Sequence Generation

To evaluate model performance on cfDNA generation, we started from real world biosamples, held out during pretraining and aimed to reconstruct them *in silico*. Two biosamples from the test set of the pretraining data were used for this analysis [64], both of which were processed using WGBS for library preparation and sequenced to a depth of 30 − 50*x*. In each sample, we designated 10% of fragments to a seed set, from which prompts were created for generation. The remaining fragments in the sample (90%) were assigned to the ground truth set. Each prompt consisted of five full fragments in the same 1kb region within the seed set and the cfDNA start special token `<cfdna>`, followed by the first three nucleotides of a fragment in the ground truth set. This target fragment was non-overlapping with the prompt and from the same 1kb region. We focused on 68 repeat-masked high-coverage 1kb regions. A full list of these regions can be found in Table S3. During generation we applied a top *k* sampling method where *k* = 2 with a temperature *T* = 0.7. Decoding terminated when either

- The cfDNA end special token `</cfdna>` was generated.
- Maximum token limit was reached, which is set to *c* − *length*(*p*), where *c* is model context (1024) and *p* is the prompt.

### 4.4 Cell Type-of-Origin Classification

#### 4.4.1 Contrastive Training Procedure and Loss

Within a marker region for any cell type, fragments from the target cell type are vastly outnumbered by fragments from other cell types. This severe class-imbalance destabilises training and can cause the model to overfit the dominant class. To counteract this problem, we adopted a contrastive learning strategy. The method clusters the scarce positive examples while repelling the overwhelming pool of negatives by (a) sampling a tractable subset of negative reads, (b) augmenting the positives, and (c) re-weighting the loss. During training, the network directly compared reads from different cell types, encouraging sequences from the same cell type to co-locate in representation space and pushing sequences from different types apart. We implemented this behaviour with a contrastive loss that combines mean-based loss and hard negative mining [111, 112]. In order to maximise model generalisability during each epoch of training, random negative examples were re-sampled for each anchor.

Mean-based loss provided stable overall training by considering an average of negative examples. Hard negative mining helped with fine-grained discrimination, especially when certain cell types share similar methylation patterns. The combination enabled the model to learn both broad distinctions and subtle differences between cell types, improving its ability to generalise.

In addition, the model performed cell type classification, where it processed methylation sequences through Pleiades base model to generate numerical embeddings and predict the corresponding cell type via a classification head. This was supervised by a classification loss, which relied upon pooled sequence representations.

To optimise both learning strategies, the model was trained with a combined loss, where each component was weighted by a multiplier (*m*_class_ and *m*_contr_) to balance their contributions.

By simultaneously learning from both tasks, the model built a deep understanding of cell type specific epigenetic features. Contrastive learning enhanced its ability to distinguish patterns, while classification provided direct supervision for accurate cell type prediction. This dual approach ensured robust representations, even in cases where some cell types were more common than others.

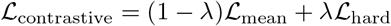

where

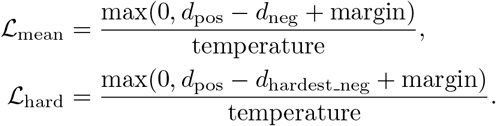

where

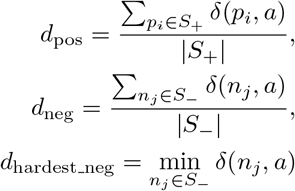

where *a* is the vector representation of the anchor sample and *S*_+_ and *S*_−_ are the set of all positive and negative example’s vector representations, respectively.

The distance metric *δ* was defined as follows:

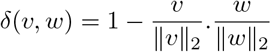

where *v* and *w* are input vectors and ∥*v*∥_2_ and ∥*w*∥_2_ represents *l*_2_ norms of these vectors. The distance metric is 1− cosine similarity.

The hyper-parameters used are:

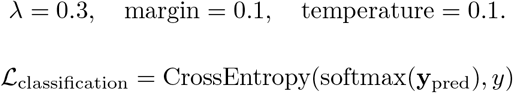

Here,

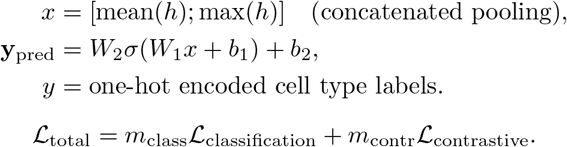

We also experimented with a regularisation term added to the contrastive loss to prevent representation collapse and increase training stability.

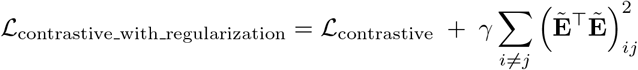

where 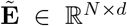 is the matrix of row-wise normalised embeddings, and *γ* is the multiplier (regularisation coefficient) controlling the strength of the diversity penalty.

#### 4.4.2 Cell-Type Differentially Methylated Regions

To fine-tune Pleiades for the CToO task, we curated a specialised dataset comprised of differentially methylated regions (DMRs) from the DNA methylation atlas [58]. To prevent information leakage into the test set, samples were first randomly split into training and test sets, and DMRs were then called exclusively from the training set. DMRs were identified for each cell type in a 1-vs-all manner to distinguish between each cell type and all-the-rest, using the open-source software wgbstools [113]. The genome segmentation and marker finding were done using `segment` and `find markers` commands with each marker region containing ≥2 CpGs, marker size from 50 to 2000bp, delta mean difference ≥0.4, significance *p*-value ≤ 0.01 and employing Bayesian pseudocounts of 15 for both C and T counts. The pseudocount acts as a regulariser that shrinks methylation estimates toward 0.5 at low-coverage CpG sites, preventing noisy observations from driving spurious block boundaries during genome segmentation. From the resulting DMRs, we selected the top 100 and top 1, 000 markers per cell type, yielding 3, 801 and 34, 134 regions in total respectively. We also used the official UXM top 25 marker set for reference [58].

For DNA sequences intersecting each DMR region, the ones belonging to the target cell type of the DMR were annotated as positives with the rest annotated as negatives. Negative reads have broadly similar methylation signals, so their labels were replaced with a generic negative label, for example `not neuron` if the DMR target is `neuron`. We used contrastive learning to force the model to separate its representations of reads for DMR targets and other cell types within each DMR region. In order to do that, each positive read within a DMR region were used as an anchor, while a constant number of other positive examples (3) are randomly selected to match with each anchor alongside a constant number of negative examples (36) from other cell types in the same region. This random sampling was done per training epoch to maximise model generalisability.

Final classification results were calculated on the total set of reads that intersected with the DMR regions. For performance comparison, we evaluated three DMR sets: the top 25 DMR set reported by [58], and Top 100 and Top 1000 DMR sets. To assess generalisation, we tested on an out-of-distribution dataset incorporating sequence data [78], using only reads intersecting the predefined DMR sets.

#### 4.4.3 Out-of-Distribution data for evaluation

For evaluating CToO and deconvolution tools, we obtained external data (Do et al., 2020) [78] and only considered healthy samples. This data set consists of 478 FACS-processed samples from various cell types. Of the total samples, 96 failed quality control criteria: 92 had low sequencing coverage and 4 showed evidence of failed bisulfite conversion. We grouped related cell types into major cell types defined by the methylation atlas [58] - six cell types were selected: B cell, monocyte & macrophage cell, T cell, liver hepatocyte, oligodendrocyte, and neuronal cells. Data was processed similarly to our cfDNA data, but with an underlying aligner such as Bismark [109] due to its native support for per-base DNA methylation calling.

The same 6 cell types were used for mixture experiments. We created 14 different cell type mixtures ranging from six pure cell type, three types of 2 cell type mix, three types of 3 cell type mix, one type of 4 cell type mix and a 6 cell type mix. Each mix had an equal proportion of cell types. The complete breakdown of the mixtures is shown in Table S4. To evaluate deconvolution sensitivity to rare cell types, we also performed a series of variable-proportion in-silico spike-in experiments. Starting from a baseline pool of B cell cfDNA fragments, we spiked in fragments from each of five additional cell types (monocyte/macrophage, T cell, hepatocyte, oligodendrocyte, and neuron) at frequencies of 0%, 0.1%, 1%, 5%, and 10%.

For a fair comparison, we used the same marker regions for both Pleiades and UXM with the corresponding cell atlas. In contrast, CelFiE required a distinct cell-atlas format and marker-region caller, and relied upon raw coverage instead of methylation percentages. Therefore we built a custom atlas for CelFiE to improve comparability of the results.

We fine-tuned Pleiades 90M over official regions from the Top 25 markers from UXM [58] and performed classification over fragments. For each sample we aggregated the predictions of all fragments and calculated the cell type composition to get a deconvolution result. The same Top 25 markers from 2.4.2 were used to run the deconvolution tool of UXM with the parameter `--rlen 2`, which is the minimum number of CpG sites in each fragment.

For CelFiE, we used scripts provided by Caggiano et al. (2021) [64] to identify DMRs for each of the 39 cell type groups, selecting the top 100 CpG sites per marker as recommended. We then estimated cell type proportions by running the authors’ deconvolution script with its default parameters and excluding any unknown cell types.

### 4.5 Neurodegenerative Disease Diagnosis

#### 4.5.1 Dataset

We create a proprietary AD cfDNA dataset. A cohort of 81 age and sex-matched patients with AD dementia and elderly healthy controls were recruited from the Cognitive Disorders Clinic at the John Radcliffe Hospital in Oxford, UK, or open day events (Fig. 3b). AD dementia patients met clinical criteria for the AD clinical syndrome, characterized by a progressive, multidomain, largely amnestic cognitive impairment. They underwent MRI and FDG-PET imaging, the results of which were in keeping with a clinical diagnosis of AD (temporo-parietal atrophy and hypometabolism). Following plasma biomarker analysis, ATN status was reviewed to confirm that these patients had a biomarker profile compatible with AD pathology, as framed by the 2018 NIA-AA research framework [83]. Elderly healthy controls were greater than 50 years old, age-matched to the AD dementia patients, and had no psychiatric or neurological illness and were not on regular psychoactive drugs. They also underwent brain MRI imaging, and only participants with a normal MRI scan, reviewed by two independent senior neurologists, were included in the study. Participants underwent in-person blood collection and face-to-face standard cognitive testing, the Addenbrooke’s Cognitive Examination-III (ACE-III), at the time of the visit. ACE scores lower than 88/100 were considered abnormal, and all healthy controls scoring below that threshold were excluded from this study. However, patients with AD were not recruited based on a fixed threshold on standard cognitive testing but rather took part in the study according to the criteria outlined above.

Blood was collected in six ethylenediaminetetraacetic acid (EDTA) tubes (10 mL each), and centrifuged (1800 g, room temperature, 10 minutes). The EDTA tubes were filled completely and gently inverted after collection to avoid coagulation. After centrifugation, plasma from all six tubes were transferred into one 50-mL polypropylene tube, mixed, aliquoted into 0.5 mL polypropylene tubes (Fluid X, Tri-coded Tube, Azenta Life Sciences), and stored at 4^°^C, until (less than 8 hours) it was transferred into a −80^°^C freezer. The time between blood collection and centrifugation was less than 30 minutes. Transfer time between 4^°^C and −80^°^C storage was *<* 20 minutes, and the samples were kept refrigerated during transport. All cryovials were anonymised, and the unique cryovial code was logged into a secure database, linked to the participant’s anonymous code and visit number.

PD samples were procured from a commercial supplier (AMSBIO). A cohort of sex- and age-matched samples were selected (Fig. 3b). Individuals with PD were defined as having the clinical features of bradykinesia with additionally rigidity and/or tremor. Healthy controls selected had no psychiatric or neurological illness and were not on regular psychoactive drugs. Samples were collected in EDTA tubes and processed similarly to AD samples.

cfDNA was extracted from 0.5-1.0ml plasma using the QIAamp Circulating Nucleic Acid Kit. Sequencing libraries were generated with the NEBNext Enzymatic Methyl-seq and sequenced on Illumina short read sequencer for depths between 30-50x. Data was processed using a modified MethylSeq pipeline [108] with BWA-Meth [114] as the aligner. Per-read methylation calls were processed using a modified MethylDackel tool [115]; non-CpG methylation calling feature was added. Processed BAM files were converted into fragments as described in Section 4.1.4.

For protein measurements, samples were shipped on dry ice to the Biomarker Factory/Fluid Biomarker Laboratory, UK Dementia Research Institute at University College London (UCL), London. The Dementia Research Institute (DRI) laboratory staff carried out the analyses. Plasma A*β*40, A*β*42, GFAP, and NfL were measured by single-molecule array (Simoa) technology using the Neurology 4-plexE assay on an HD-X analyzer (Quanterix), according to manufacturer’s instructions. Plasma p-Tau181 and p-Tau217 were also measured by Simoa using the pTau-181 Advantage and ALZpath assays on an HD-X analyzer (Quanterix). Samples were analysed in one round of experiments using one batch of reagents with intra-assay coefficients of variation below 10% and the analysts blinded to clinical data.

Since using the entire human genome space would be computationally prohibitive, we utilised cell type specific DMRs to extract regions for microglia, neuron, B cells, and T cells. We utilised the top 100 DMRs as described in Section 4.4.2 for all but microglia, because they are not included in the methylation atlas. For the latter, we used the 59 DMRs proposed by Tian et al. (2023) [76].

#### 4.5.2 Marker Discovery Methodology

We addressed the challenges of discovering markers for neurodegenerative diseases, such as the vast exploration space, low signal-to-noise ratio, and signal heterogeneity by employing Pleiades to search the epigenome, amplify weak signals, and capture cell-type-specific patterns. We curated data based upon biological relevance, targeting specific cell-type DMRs.

Starting from 59 DMRs for microglia and 100 DMRs each for neurons, B cells and T cells, Pleiades was utilised to further filter them to a concise set of high-confidence candidates, using nested 5-fold cross validation. The inner CV folds were used to train the region level HAT models and select the best marker regions, while outer folds were used for final fine-tuning and performance measurement. The overall workflow is summarised in Table 4.

**Table 4:** Simplified Three-Step Workflow for Diagnostic Marker Discovery.

|  | Marker Filtering | Region-Level Training | Sample-Level Training |
| --- | --- | --- | --- |
| **Goal** | Find regions that carry useful signals | Learn patterns from regions using weak labels | Make final predictions for each sample |
| **Supervision** | Weak supervision | Weak supervision | Full supervision |
| **Model Type** | 1<sup>st</sup> order HAT | 1<sup>st</sup> order HAT | 2<sup>nd</sup> order HAT |
| **Approach** | Use cross-validation to select the most informative regions based on performance and statistics | Train models on region data using labels from whole samples; represent each region with embeddings | Train a final model using true sample labels and region-level outputs |
| **Input Data** | All genomic regions | Filtered regions from previous step | Same filtered regions |
| **Output** | Selected genomic regions as biomarkers | Region-level predictions or embeddings | Final diagnosis for each sample |

To account for low signal-to-noise ratio and high variability, we selected regions with AUROC*>* 0.6 for Alzheimer’s disease and AUROC*>* 0.65 for Parkinson’s disease as well as significance of *p*-value *<* 0.01 for at least four of five inner folds across the last 50 epochs of training. Each inner CV fold was trained for 100 epochs, and validation AUROC scores from the last 50 epochs were examined. We applied a Student’s t-test to assess statistical significance above the threshold. Regions passing the t-test in at least four of the five inner CV folds were sorted by the number of passing folds and average AUROC score in descending order. The top four were selected for further training phases to fit within GPU memory constraints, particularly for larger models.

Pleiades hierarchical architecture leverages the natural biological structure of cfDNA fragmentomics, progressing from the fragment level to the sample level. It builds on a base sequence-level model by introducing a region-level model. This region-level model is fine-tuned on the final, refined set of regions, after the inner cross validation step. Finally, a sample-level model is trained to predict the Alzheimer’s disease condition. This three-tiered hierarchy — fragment → region → sample — allows the model to learn structured, set-based features in a computationally efficient manner. Importantly, it avoids the need for long-context models, which are constrained by the quadratic complexity of standard Transformer architectures. The negative control experiment using spike-in DNA (pUC19 and Lambda; Fig. S10b) confirmed that the selection procedure rejects regions lacking biological signal, providing further evidence against overfitting to technical artefacts.

#### 4.5.3 Learning Biological Hierarchical Representations

##### Fragment Level Representation Learning

We fine-tuned the base Pleiades model to encapsulate representations of cfDNA fragments into a single `[CLS]` token. A reconstruction head is used to train the base model to pack entire sequence representations in the `[CLS]` token. The intuition is that, if the `[CLS]` token can support accurate reconstruction, then it must preserve rich information about the original cfDNA fragment. The reconstruction post training is done prior to any downstream tasks finetuning and the base model is frozen after that. Table 5 shows the details of the reconstruction head architecture. It is a lightweight transformer decoder which reconstructs the input sequence conditioned on `[CLS]` token sequence embeddings. It functions as an autoencoder, incorporating global context from the `[CLS]` token via cross-attention. With two layers, it remains computationally efficient compared to the base model. Dropout was applied to attention and feedforward layers to mitigate over-fitting. To promote diverse and informative representations, we combined a diversity loss with the reconstruction loss, penalising similarity among sequence embeddings and preventing representation collapse. This dual objective captures input content and structural properties in a compact embedding space.

**Table 5:**
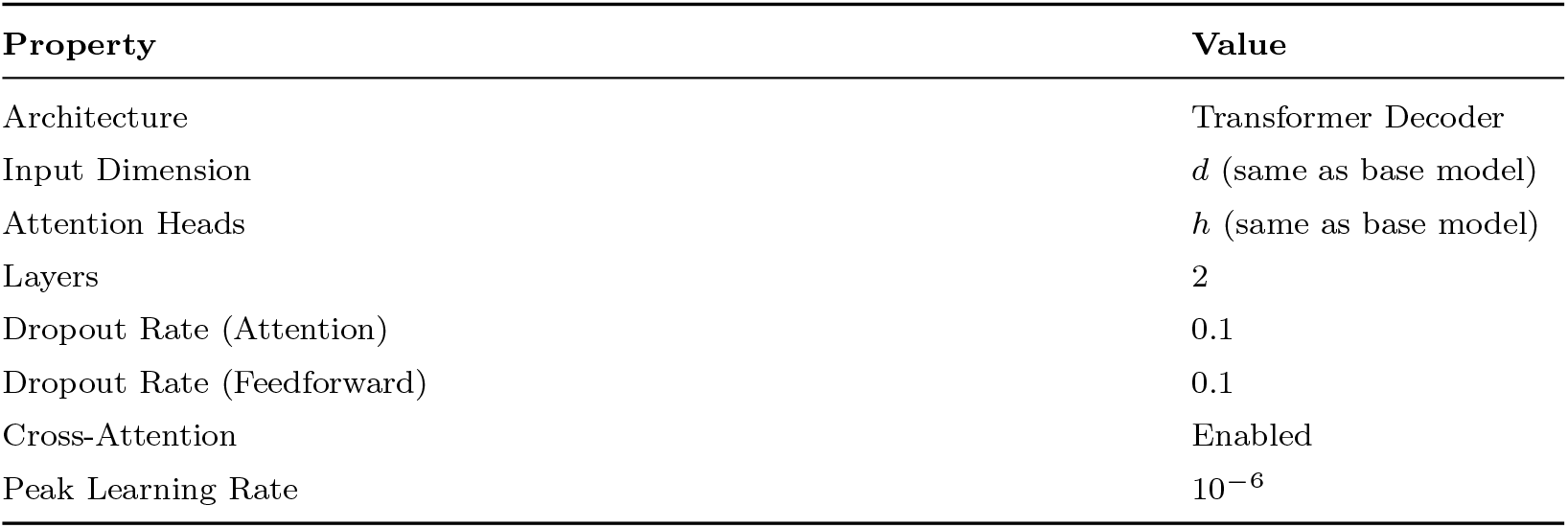
Reconstruction Head Configuration.

We defined the reconstruction head’s input and output mathematically as follows. The input sequence is given by:

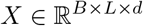

where *B* is the batch size, *L* is the sequence length, and *d* is the embedding dimension. Sequence embeddings are computed by extracting the `[CLS]` token representation from the input and used to form the context:

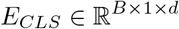

The decoder receives both the input sequence and the context embeddings, producing token-level output logits:

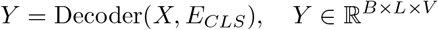

where *V* is the vocabulary size.

Training was supervised via a combination of two loss functions. The first is the standard reconstruction loss:

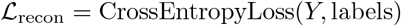

which encourages the model to accurately reconstruct the input sequence. The second is a correlation-based diversity loss:

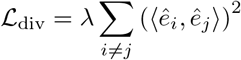

where 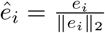 is the normalised `[CLS]` embedding for sample *i*, and *λ* is a small scaling factor, in our case 10^−6^. This loss penalises cosine similarity between distinct embeddings, encouraging orthogonality and diversity across the batch. The total loss is the sum of both components:

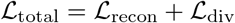

This setup enabled the model to learn compact, informative embeddings that serve downstream tasks while remaining efficient and robust.

##### Region Level Representation Learning

By keeping the base model weights frozen, we trained a hierarchical transformer with a classification head on the task of predicting whether an individual has the target disease. The input to the model is a batch of fragments grouped in genomic regions. The base model outputs the embedding representations of these fragments, which are grouped accordingly to form a region. This collection of fragment embeddings is the input to the second transformer, which is trained in a weakly supervised way; the disease label is assigned to each of the regions of the sample’s DNA. The model is guided by a classification head that learns to predict these labels.

The input to the base model is *X* ∈ ℝ^*B×L×d*^ and the sequence embeddings are described by *E*_*CLS*_ ∈ ℝ^*B×d*^

The architecture groups sequences by their region IDs. Let

- **R** ∈ ℕ^*B*^ denote the region ID assigned to each of the *B* sequences,
- **R**_unique_ = unique(**R**) be the set of unique region IDs,
- *N* = |**R**_unique_| be the total number of distinct regions.

For each region *r* ∈ **R**_unique_, define the set of embeddings associated with that region as:

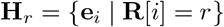

Here, 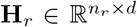, where *n*_*r*_ is the number of sequences belonging to region *r*, and *d* is the embedding dimension.

Each region-specific embedding set **H**_*r*_ is then passed through an encoder:

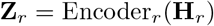

which is used by a classification head *f*_*r*_ with the following loss function:

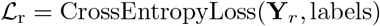

where **Y**_*r*_ = *f*_*r*_(**Z**_*r*_). To encourage diversity among attention heads by penalising their similarity, we define the loss ℒ_attndiv_ as:

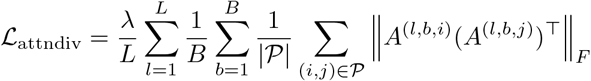

where:

- *λ*: scaling factor,
- *A*^(*l,b,h*)^ ∈ ℝ^*S×S*^ : normalised attention map,
- *P* : all head pairs (*i, j*), *i < j*,
- ∥ · ∥_*F*_ : Frobenius norm,
- *L, B, H, S*: number of layers, batches, heads, and sequence length.

The total loss becomes:

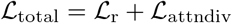

##### Sample Level Representation Learning

The architecture can be extended hierarchically by reusing the base and the region level models and adding a third sample-level model. The first two models were then frozen and the third received its inputs by grouping region-level embeddings into individual samples. Let:

- **S** ∈ ℕ^*N*^ denote the sample ID assigned to each of the *N* regions,
- **S**_unique_ = unique(**S**) be the set of unique sample IDs,
- *M* = |**S**_unique_| be the total number of distinct samples.

For each sample *s* ∈ **S**_unique_, we collected the embeddings of all regions belonging to that sample:

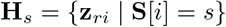

Here, 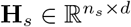, where *n*_*s*_ is the number of regions in sample *s* and *r* ∈ **R**_unique_.

Each sample-level set of region embeddings **H**_*s*_ was then passed through a transformer encoder:

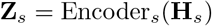

and the sample representation **z**_*s*_ ∈ ℝ^*d*^ was passed to a classification head *f*_*s*_.

The model was trained by assigning the sample’s condition label (e.g., AD vs. non-AD) to the collection of all regions. The classification loss was defined as:

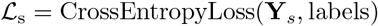

where **Y**_*s*_ = *f*_*s*_(**Z**_*s*_).

The overall loss combined attention diversity regularisation, and sample-level classification:

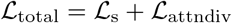

#### 4.5.4 Baseline diagnosis classifiers

##### DELFI

Following the methodology of Cristiano et al. [63], we computed fragment length ratio features from cfDNA EM-Seq data in 5 Mb genome-wide bins, extracting the ratio of short fragments (below a 151 bp threshold) and the corrected fragment count per bin. Features from sex chromosomes were removed, and the remaining features were standardised (zero mean, unit variance) using statistics computed on the training set. We trained an L1/L2-regularised logistic regression classifier using 5-fold nested cross-validation, with the regularisation strength C and penalty type selected by grid search over inner folds.

##### Combined fragmentomics + methylation linear model (Meth + Length + End-motif)

As a non-foundation-model comparator, and inspired by the methylation-classifier approach in Caggiano et al. [90], we trained a single logistic-regression classifier on conventional cfDNA features extracted from the same per-cell-type top-100 differentially methylated regions (DMRs) used by Pleiades (B cells, T cells, microglia, and neurons). For each sample we pooled all cfDNA fragments overlapping the DMRs in a region set and computed three feature families: (i) context-specific methylation rates (CpG, CHG, CHH) from the per-read Bismark methylation calls; (ii) the fragment-length distribution binned into nine cfDNA-informed length intervals (edges at 100/130/150/167/180/200/250/300 bp) and normalised to fractions; and (iii) 5^*′*^ end-motif frequencies, with the motif resolution chosen by a sweep over *k* = 1–4 (*k* = 1, the four 5^*′*^-terminal nucleotide frequencies, was optimal), giving a 16-dimensional feature vector. Features were standardised on the training fold and classified with L2-regularised logistic regression (*C* = 0.5, balanced class weights). For each cell type, features were aggregated over that cell type’s DMRs; the “All Regions” model pooled fragments across all studied cell types. A gradient-boosted-tree variant performed comparably; we report the linear model for interpretability.

##### MethylGPT

We fine-tuned MethylGPT [91], a pretrained transformer model originally trained on Illumina 450K array methylation data. Following the authors’ recommended fine-tuning procedure, beta values at 49,156 Illumina probe positions were extracted from genome-wide wgbstools [113] .beta files (minimum 3 reads per CpG), and the pretrained encoder was frozen while a two-layer MLP classification head was trained with BCE loss for 50 epochs using AdamW (lr = 110^−3^).

##### CpGPT

We fine-tuned CpGPT [16], a 2.9M-parameter DNA methylation foundation model. Although CpGPT accepts arbitrary genomic CpG positions, following the approach used in the original publication we extracted beta values at Illumina array CpG positions from genome-wide wgbstools [113] .beta files and formatted them into the model’s native memory-mapped input representation. CpG sites were retained if covered by at least 3 reads in 100% of AD samples or 95% of PD samples (due to the larger size of the PD cohort). The pretrained model was fine-tuned end-to-end using CpGPT’s built-in condition decoder head with its default weighted loss combination and AdamW ScheduleFree optimiser (lr = 1 × 10^−4^) for 50 epochs.

#### 4.5.5 Evaluation

The performance of Pleiades models, baseline classifiers, and protein biomarkers is measured using AUROC metric on the outer 5-fold cross validation of both our disease datasets. All results were reported as mean of the 5 outer folds in Fig. 3 and Fig. S10. Statistical significance was assessed using one-sided paired *t*-tests on the per-fold AUROCs across the five outer cross-validation folds, comparing model sizes within each cell type and the combined Pleiades 7B + pTau-217 model against each proteomic marker, with Benjamini–Hochberg correction for multiple comparisons (Fig. 3c-e; Supp. Table S5).

#### 4.5.6 Gene Ontology Enrichment Analysis

To assess the biological relevance of the HAT-selected marker regions, we performed Gene Ontology (GO) enrichment analysis using GREAT (Genomic Regions Enrichment of Annotations Tool) [84] via the rGREAT R package [85]. The unique top-4 regions pooled across all four cell types were tested for enrichment of GO Biological Process, Molecular Function, and Cellular Component terms against the full set of starting candidate DMRs as background, ensuring that enrichment reflects the model’s selection rather than properties of the candidate region set. GREAT assigns biological annotations to genomic regions based on nearby genes using regulatory domain assignment and tests enrichment with a binomial test over genome fraction. P-values were corrected for multiple testing using the Benjamini–Hochberg method across all 724 tested GO terms.

### 4.6 Mechanistic Interpretability

All mechanistic-interpretability analyses were performed on the final-layer `CLS`-token embeddings of Pleiades 7B. Embeddings were taken from the output of the last transformer block (layer 42), before the final normalisation layer, for both the reconstruction-tuned and pretrained-only checkpoints.

#### 4.6.1 Principal Component Analysis Visualisation

To visualise how Pleiades organises fragment-length information, we pooled `CLS` embeddings across the four cell types, restricted to the top-4 regions per cell type; that is, to the genomic regions actually selected by the 2nd order HAT classifier during fine-tuning. For *N* sample data points, we then fit principal component analysis on the resulting *N* × 4,096 pooled embedding matrix, retaining the first 10 components, and visualised the first two and three components coloured by fragment length. PCA was computed for the reconstruction-tuned 7B activations (Fig. S12a).

#### 4.6.2 Sparse Autoencoders

##### Training Data

For sparse autoencoder (SAE) training, we use the same subset of cfDNA pretraining dataset as above for linear probing.

###### SAE Hyperparameters

We trained a Top-*K* sparse autoencoder [92] on the SAE training data. Let *x* ∈ ℝ^*d*^ denote a `CLS` activation with *d* = 4,096. The SAE comprises an encoder with weights 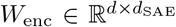 and bias 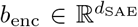; a decoder with weights 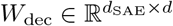 and bias *b*_dec_ ∈ ℝ^*d*^, which also serves as a pre-encoder offset following [92]; and a TopK(·) operator that retains the *K* largest entries of its argument and sets the remainder to zero. The sparse code 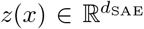 (with at most *K* nonzero entries) and the reconstruction 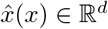 are

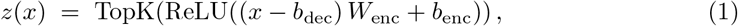

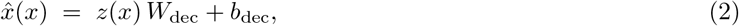

with latent dimension *d*_SAE_ = 4*d* = 16,384 (expansion factor 4) and *K* = 64 active features per input. The rows of *W*_dec_, which are the decoder dictionary atoms, one per latent feature, were normalised to unit *ℓ*_2_-norm after every optimiser step.

The training objective combined a mean-squared reconstruction loss, 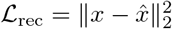, with the auxiliary dead-feature loss of Gao *et al*. [92], which re-engages features whose cumulative activation count has fallen below a threshold (10^7^ tokens) by training them to reconstruct the residual 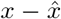 using only their own top-*K* contribution; this auxiliary term was weighted by 1*/*32. Optimisation used Adam (*β*_1_ = 0.9, *β*_2_ = 0.999) with a peak learning rate of 3 × 10^−4^, 250 steps of linear warmup followed by cosine decay, a batch size of 16,384, and mixed-precision (bfloat16) matrix multiplications with fp32 loss accumulation. Five percent of the pretraining activations were held out as a validation set. Training was terminated by early stopping with a patience of 5 evaluations on validation reconstruction MSE, and the best validation checkpoint was used for all downstream analyses. Of the *d*_SAE_ = 16,384 features, 761 (4.64%) fired on at least one fragment of the AD-DMR matched subsample; the remaining 15,623 dead features were dropped prior to all downstream analyses.

###### Hierarchical Set Model → SAE Gradient Attribution

We quantified the contribution of each SAE feature to the AD classifier’s decisions by attributing its output logit back through the SAE. Attribution was computed for every combination of cell type and outer cross-validation fold, yielding 20 runs. In each run we loaded the corresponding fold specific sample level model checkpoint and applied it to the full 81-patient AD cohort, restricted to the model’s selected regions. Fragments from each (patient, region) pair were passed through the reconstruction-tuned Pleiades 7B to obtain `CLS` activations *x*, which were encoded through the trained SAE to yield sparse feature vectors *z*(*x*). The sample model classifier was then evaluated on 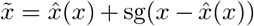, where sg denotes a stop-gradient; this leaves the forward pass unchanged while routing gradients with respect to *z* exclusively through the SAE reconstruction path. All sample model parameters were frozen.

For each fragment *f* and feature *j*, the attribution was computed as

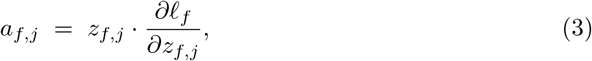

where *ℓ*_*f*_ is the logit of the predicted class for the patient to which *f* belongs; this product of activation and gradient is equivalent to a single-point Taylor attribution [116]. Within each (cell type, fold), each feature’s attributions were summed over fragments and normalised so they sum to 100% before averaging across the 20 experiments. Feature-level importance for a given (cell type, fold) was summarised as the percent attribution

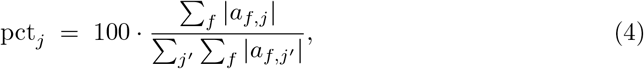

and Fig. 4a reports the mean ± s.d. of pct_*j*_ across the 20 (cell type × fold) runs for the 20 highest-ranked features. Tuning curves (Fig. 4e) were constructed from an independent stratified subsample of 10,000 fragments per cell type, plotting each top-ranked feature’s sparse activation against cfDNA fragment length.

###### Mutual Information Estimation

Mutual information between SAE feature activations and per-fragment annotations was estimated using scikit-learn’s `mutual info regression` and `mutual info classif` functions for continuous and discrete targets respectively, with `n neighbors` = 3 [117].

###### Top-Feature Enrichment Test

To test whether the top-20 attribution-ranked features carried more of each biological signal than expected by chance, we compared their mean mutual information against a null distribution of 1,000 random draws of 20 features sampled without replacement from the alive features outside the top 20. For each signal (fragment length, CpG and CpH methylation ratio, and genomic region identity), the one-sided empirical *p*-value was the fraction of null draws whose mean mutual information met or exceeded the observed top-20 mean, and these *p*-values were Benjamini–Hochberg corrected across the four signals to yield the reported *q*-values (Fig. 4b).

#### 4.6.3 SAE Feature Steering

Steering was implemented as a per fragment, per region edit to the L42 `CLS`-token activation, designed to isolate one SAE feature’s causal contribution to AD classification while keeping the rest of each patient’s input unchanged. For each (cell type, fold) we used the matching fold trained sample-level model together with that fold’s selected regions. For a target SAE feature *f* , steering multiplier *m*, target region *d*, and `CLS`-token activation *x* of a fragment within that patient’s input, we have:

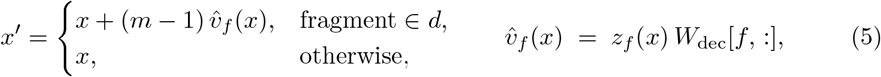

where 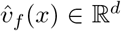 is feature *f* ‘s contribution to the SAE reconstruction of *x*; the product of *f* ‘s scalar activation *z*_*f*_ (*x*) on that fragment and the decoder dictionary atom *W*_dec_[*f*, :]. Setting *m* = 0 subtracts the feature’s contribution in full (zero ablation), *m* = 1 recovers the unmodified baseline, and *m >* 1 amplifies the feature in proportion to its native activation. The multiplier was swept on an 18-point grid covering [0, 30]: half-steps in [0.0, 3.0] near baseline, unit steps in [3.0, 10.0], and 5-unit steps from 15 to 30 to probe far amplification saturation. For each (patient, *d, f, m*) the steered *x*^*′*^ vectors were passed through the model’s final norm and pooling, the bottleneck, the HAT, and the classification head of the fold matched sample model, and *P* (AD) was taken as the softmax probability of the AD class. Aggregation proceeded in two stages: first averaging *P* (AD) across all scored patients within a fold (cases and controls pooled, as the steering effect is defined on each patient relative to its own baseline), then averaging across the folds in which the region was selected. The 20 SAE features steered are a single global set: the top-20 features by mean HAT `pct`-attribution across all 20 (cell type × fold) sample models. Consequently “feature rank *k*” denotes the same SAE feature index across all cell types and all folds, and each cell-type figure (Fig. S13) varies only in the downstream sample model used to score the steered activations. The *m* = 1 value was subtracted from each multiplier to yield Δ*P* (AD). Mean |Δlogit| was computed as the per patient absolute shift in the AD logit at *m* = 0, averaged over scored patients and then over folds, matching Fig. 4a.

#### 4.6.4 Fragment Length Classifiers

Three fragment length baselines were evaluated alongside the sample level model. Each is a logistic regression with *L*_2_ regularisation (*C* = 1.0, LBFGS, up to 2000 iterations) on per region median fragment length, fit on the fold’s training patients and scored on the fold’s held-out test patients, matching the sample model’s data split exactly. Features were *z*-scored using statistics computed on the training split and applied to the test split. The three baselines differ only in their choice of regions:

##### Frag. Length (Pleiades regions)

Each patient was described by four features representing the median fragment length over fragments overlapping one of the regions that Pleiades had originally selected.

##### Frag. Length (random regions)

The design as above, but with the four attribution-selected regions replaced by four regions drawn uniformly at random. To stabilize the estimate, 20 independent random draws were performed per (cell type, fold), giving 20 AUROCs per fold which were averaged into a single per-fold random-control AUROC prior to the cross-model statistical test (Table S6).

##### DELFI 151bp

The published DELFI fragmentation-based ctDNA classifier evaluated at the 151 bp short-fragment cutoff [63].

## 5 Data Availability

Public datasets used in this study are available from their original sources as cited: the DNA methylation atlas of normal human cell types [58], cfDNA whole-genome bisulfite sequencing from healthy individuals [64], the 1000 Genomes Project reference [65], and the Nucleotide Transformer benchmark [15]. The clinical cfDNA datasets generated in this study (Alzheimer’s disease, Parkinson’s disease, and the in-house healthy cfDNA used for pretraining) are subject to participant consent and ethical approval restrictions that preclude open public release. Limited access for non-commercial academic research will be provided to qualified researchers upon publication, through controlled-access APIs consistent with the terms under which the data were originally collected and subject to appropriate data-sharing and ethics agreements. The specific access procedure will be agreed across contributing institutions by the proof stage.

## 6 Code Availability

A summary repository is hosted at https://github.com/primamente/pleiades. The analysis code and evaluation pipelines associated with this study will be released for non-commercial academic research use upon publication.

## 7 Model Availability

Pleiades models (90M, 600M, and 7B parameters) will be made available for non-commercial academic research use upon publication, through controlled-access APIs operated by Prima Mente. Access will be granted to qualified researchers on terms consistent with the model’s research-only license.

## 8 Acknowledgements

We thank Vivek Natarajan, Eric Nguyen, Yuval Dor, and Taya Reed for their support and contributions to this work. We are also grateful to the wider Prima Mente team, including Marie Schildt, Devin Gilliam, and Hayley Holt. This project was supported in great part by resources and services provided by NVIDIA DGX Lepton, Nebius, Siam AI, Eternis Labs, and Google Cloud Platform.

## 9 Author Contributions

P.N. and C.N. co-led the design and architecture of the base model. J.-O.G., P.N., and N.L. created and contributed to the data corpus. P.N., C.N., J.-O.G., D.By. and P.K. did the main experiments. C.F., N.K.W., M.B., D.H., M.T.P., A.Ja. and D.Ba. worked on mechanistic interpretability experiments. H.B. and A.Jh. performed the wet lab processes to generate the main diagnosis datasets. S.To., M.H., S.M., and S.Th. performed clinical assessment of patient cohorts utilised in this study, and contributed biosamples for experimental analysis. A.W., A.Kar. and A.Kam. contributed to experiments. L.G., W.R., T.L., R.Su. and A.M. contributed software that helped with experiments. F.M.M.Z., K.S., H.M., I.K., H.Z., J.C.M.W. and R.So. contributed to experiment design and clinical direction. R.So. and P.N. conceived of and co-led the project.

## 10 Declarations

Husam Babikir, Donal Byrne, Javkhlan-Ochir Ganbat, Luca Giacomoni, Anjeet Jhutty, Arihant Kamdar, Andrey Karailiev, Pooja Kathail, Timing Liu, Hannah Madan, Francisco M Martín-Zamora, Christoforos Nalmpantis, Pouya Niki, Will Rowe, Ravi Solanki, Robert Sugar, Jonathan C. M. Wan and Alfred Wong are shareholders of Prima Mente.

Henrik Zetterberg (HK) is a Wallenberg Scholar and a Distinguished Professor at the Swedish Research Council supported by grants from the Swedish Research Council (#2023-00356, #2022-01018 and #2019-02397), the European Union’s Horizon Europe research and innovation programme under grant agreement No 101053962, and Swedish State Support for Clinical Research (#ALFGBG-71320). The UK DRI Biomarker Factory is funded by the National Institute for Health and Care Research University College London Hospitals Biomedical Research Centre, the UK Dementia Research Institute at UCL (UKDRI-1003), and the Weston Family Foundation. HZ has served at scientific advisory boards and/or as a consultant for Abbvie, Acumen, Alector, Alzinova, ALZpath, Amylyx, Annexon, Apellis, Artery Therapeutics, AZTherapies, Cognito Therapeutics, CogRx, Denali, Eisai, Enigma, LabCorp, Merck Sharp & Dohme, Merry Life, Nervgen, Novo Nordisk, Optoceutics, Passage Bio, Pinteon Therapeutics, Prothena, Quanterix, Red Abbey Labs, reMYND, Roche, Samumed, ScandiBio Therapeutics AB, Siemens Healthineers, Triplet Therapeutics, and Wave, has given lectures sponsored by Alzecure, BioArctic, Biogen, Cellectricon, Fujirebio, LabCorp, Lilly, Novo Nordisk, Oy Medix Biochemica AB, Roche, and WebMD, is a cofounder of Brain Biomarker Solutions in Gothenburg AB (BBS), which is a part of the GU Ventures Incubator Program, and is a shareholder of Prima Mente and MicThera (outside submitted work).

Ivan Koychev (IK) has received honoraria for advisory board roles from J&J and Novo Nordisk and non-promotional speaker fees from Eisai, is in receipt of an investigator-initiated grant from Novo Nordisk to explore the effects of a GLP-1 receptor agonist in preclinical dementia and is a medical advisor (stock options and/or retainer fees) to the following health technology companies: Five Lives, Oxford Brain Diagnostics, Leaf AI, Paloma Health and Prima Mente.

Sofia Toniolo (ST), Masud Husain (MH), Sian Thompson (SiT) are funded by the Wellcome Trust. Sanjay G. Manohar (SGM) is funded by a Medical Research Council (MRC) Clinician Scientist Fellowship and National Institute of Health and Care Research (NIHR) Oxford Biomedical Research Centre (BRC) and NIHR Oxford Health BRC. MH has received speaker and advisory board honoraria from Lilly, Otsuka, and Sumitomo.

Khaled Saab is a shareholder in Alphabet, Inc.

Netanel Loyfer, ST, SiT, and SGM declare no conflict of interest.

Two patents have been filed encompassing this work.

