## Supplementary Information for "Human whole-epigenome modelling for clinical applications with Pleiades"

### Supplementary Materials

#### S1: Nucleotide Transformer Benchmarks

The official Nucleotide Transformer Benchmarks [15] consist of 18 classification tasks, where the sequences with the positive label are sampled from genomic regions with the special characteristics such as promoter and enhancer, and the negative sequences are sampled from the rest of the genome. Negative sequences in the official benchmark are not fully random, with the genomic positions of the random sequences starting only from genomic loci divisible by 1000. Fig. S2a depicts this bias in genomic start positions. Sequences with label 0, always start from genomic loci that have modulo 1000 with respect to 1000, but sequences with label 1 and 2 have a uniformly random distribution with respect to start position modulo 1000.

This systematic offset introduces a positional bias that inadvertently reduces the diversity of the negative set. Because Pleiades explicitly encodes genomic coordinates, it achieves near-perfect scores on the original NT benchmarks (see Fig. S2c)

In order to fix this bias, we add a random jitter in the range of $[-500, 499)$ to the start positions of negative sequences (Fig. S2b). All three labels have an indistinguishable distribution with respect to the start position modulo 1000. We called this new dataset the *Unbiased* Nucleotide Transformer Benchmark.

Fig. S2d shows that our results after fine-tuning the baseline models DNABERT-2 and NT MS 2.5B for five epochs on our Unbiased NT Benchmarks.

**Table S1**: Performance (Matthews Correlation Coefficient, MCC) of all models on unbiased Nucleotide-Transformer benchmark tasks (values rounded to four decimal places).

| Task | DNABERT-2 | NT MS 2.5 B | Pleiades 90 M | Pleiades 600 M | Pleiades 7 B |
|---|---|---|---|---|---|
| H2AFZ | 0.4903 | 0.5358 | 0.7325 | 0.7147 | **0.9837** |
| H3K27ac | 0.4778 | 0.5141 | 0.7658 | 0.7390 | **0.9988** |
| H3K27me3 | 0.5965 | 0.6294 | 0.7893 | 0.7948 | **0.9912** |
| H3K36me3 | 0.6387 | 0.6636 | 0.7391 | 0.8043 | **0.9953** |
| H3K4me1 | 0.4353 | 0.4886 | 0.7602 | 0.7466 | **0.9933** |
| H3K4me2 | 0.5523 | 0.5695 | 0.7651 | 0.7661 | **0.9915** |
| H3K4me3 | 0.6171 | 0.6328 | 0.7148 | 0.6827 | **0.9768** |
| H3K9ac | 0.5662 | 0.5423 | 0.7307 | 0.7001 | **0.9960** |
| H3K9me3 | 0.4675 | 0.4876 | 0.6996 | 0.7063 | **0.9670** |
| H4K20me1 | 0.6434 | 0.6608 | 0.8192 | 0.8051 | **0.9982** |
| Enhancers | 0.5231 | 0.5461 | 0.6230 | 0.6297 | **0.9770** |
| Enhancers (types) | 0.4658 | 0.5031 | 0.5860 | 0.5977 | **0.9000** |
| Promoters | 0.7546 | 0.7581 | 0.7416 | 0.7540 | **0.9759** |
| Promoters (non-TATA) | 0.7440 | 0.7921 | 0.7530 | 0.7643 | **0.9725** |
| Promoters (TATA) | 0.7767 | 0.8211 | 0.6002 | 0.7616 | **0.9908** |
| Splicing (Acceptors) | 0.8308 | **0.9657** | 0.9509 | 0.9593 | 0.9644 |
| Splicing (All) | 0.8496 | **0.9693** | 0.9597 | 0.9481 | 0.9659 |
| Splicing (Donors) | 0.8309 | **0.9736** | 0.9652 | 0.9694 | 0.9660 |

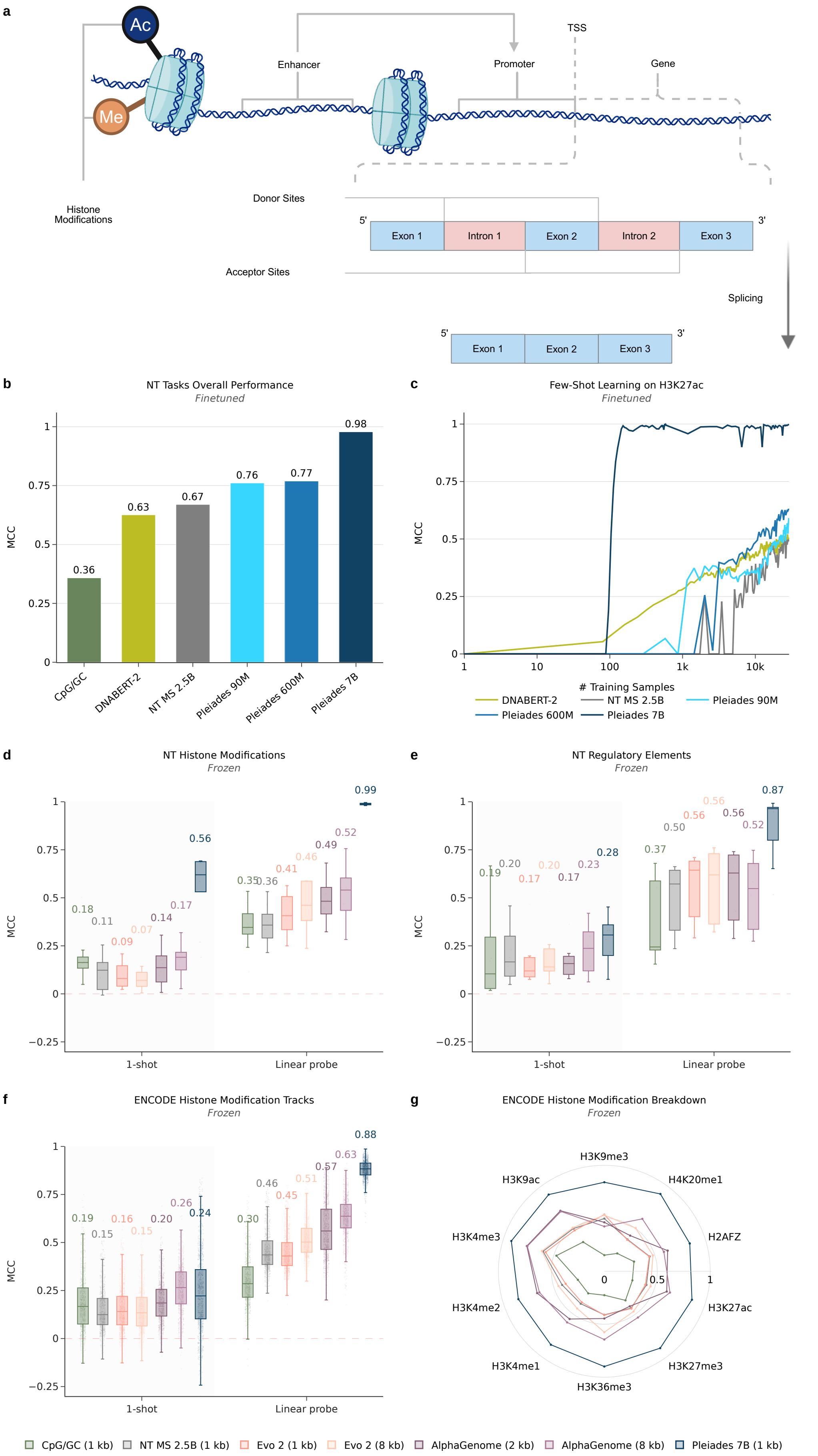


**Fig. S1**: **Nucleotide Transformer and ENCODE benchmarking of Pleiades (extended).** **(a)** Schematic of the Nucleotide Transformer (NT) benchmark task families. **(b)** Overall fine-tuned performance across the 18 NT tasks, shown as the mean Matthews Correlation Coefficient (MCC) per model (CpG/GC, DNABERT-2, NT MS 2.5B, Pleiades 90M, 600M and 7B). **(c)** Few-shot learning on the H3K27ac histone-modification task: fine-tuned MCC as a function of the number of training sequences (log scale) for each model (DNABERT-2, NT MS 2.5B, Pleiades [illegible]

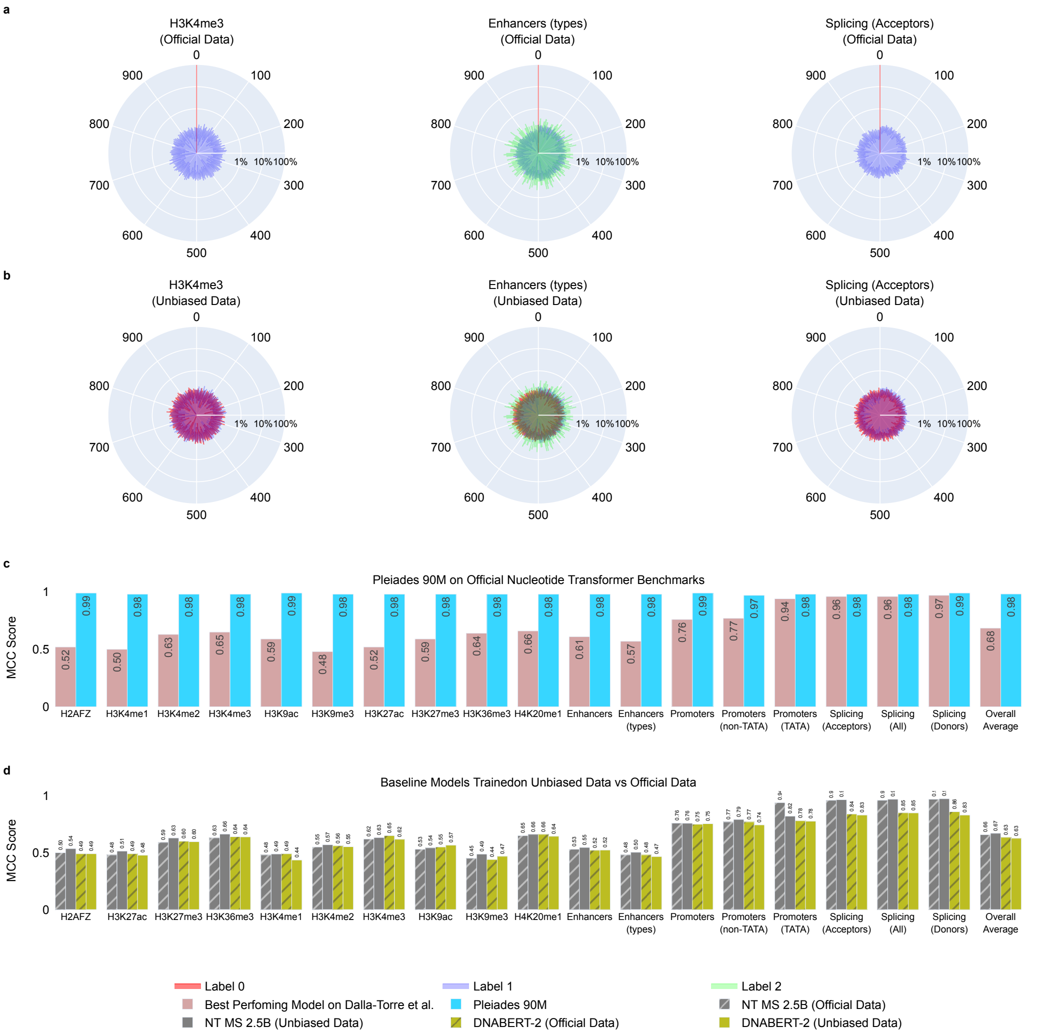


**Fig. S2**: **NT Benchmarks Positional Bias, Unbiased Dataset and Performance Comparisons** **(a)** Polar plot showing distribution of start position modulo 1000 of Official NT Benchmarks, separated by label. **(b)** Unbiased dataset where the bias is removed. **(c)** Comparison of MCC for Pleiades 90M vs Best Performing Baseline Models from Nucleotide Transformer [15] on official NT Benchmarks Data. **(d)** MCC Comparison for Baseline Models on NT-Benchmarks Official Data vs our Unbiased Data.

**Table S2: CpG-shortcut analysis on the unbiased NT histone-mark benchmarks.** All metrics are computed on raw DNA sequence inputs and aggregated over the 9 histone marks whose positive class concentrates at high-CpG loci (H3K9me3 excluded, as its inverted CpG–label relationship would flip the sign of the bias columns and render them non-comparable). The CpG-only baseline is a parameter-free classifier that thresholds per-locus CpG density at the per-mark median. The FPR bias is computed as FPR(CpG rich) - FPR(CpG poor) and the FNR bias is computed as FNR(CpG poor) - FNR(CpG rich). Higher bias indicates a greater over-reliance on CpG density as a decision shortcut in histone mark prediction.

| | | Δ vs. | FPR | | | FNR | | |
|---|---|---|---|---|---|---|---|---|
| Model | Mean MCC | CpG-only | CpG rich | CpG poor | bias | CpG rich | CpG poor | bias |
| CpG-only baseline | 0.310 | — | — | — | — | — | — | — |
| DNABERT-2 | 0.579 | +0.27 | 0.481 | 0.184 | +0.297 | 0.072 | 0.297 | +0.225 |
| NT MS 2.5B | 0.589 | +0.28 | 0.497 | 0.201 | +0.296 | 0.062 | 0.250 | +0.189 |
| Pleiades 90M | 0.763 | +0.45 | 0.209 | 0.074 | +0.135 | 0.077 | 0.206 | +0.129 |
| Pleiades 600M | 0.752 | +0.44 | 0.237 | 0.072 | +0.165 | 0.065 | 0.240 | +0.175 |
| **Pleiades 7B** | **0.994** | **+0.68** | **0.002** | **0.002** | **+0.000** | **0.004** | **0.002** | **−0.001** |

### S2: cfDNA Generation

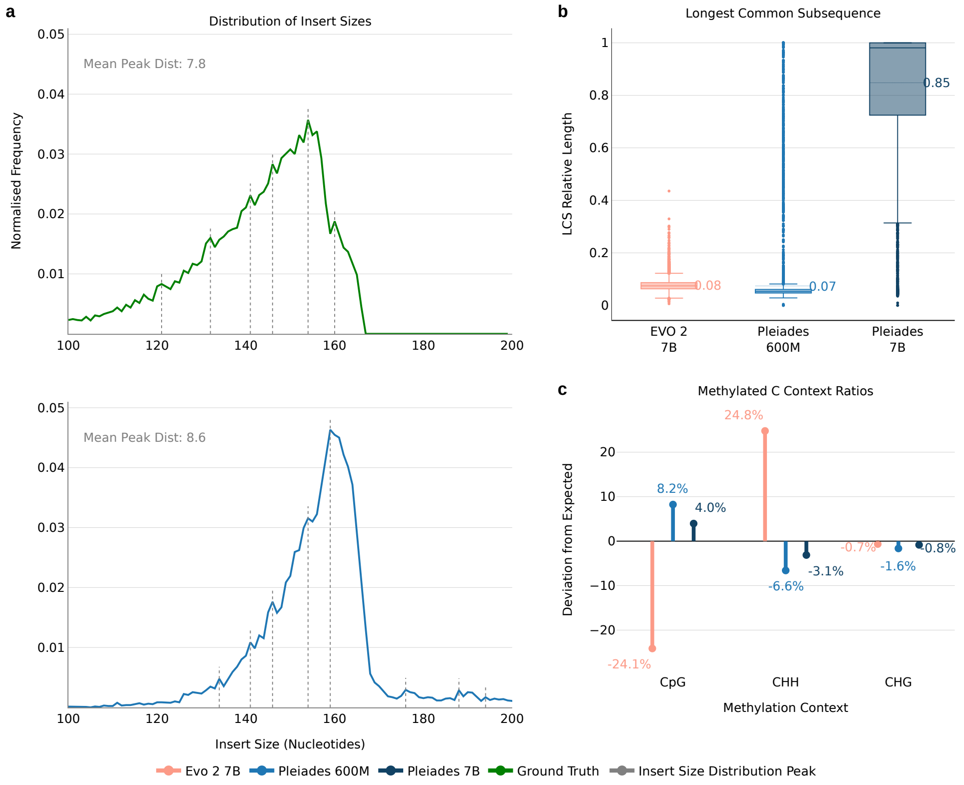


**Fig. S3**: ***In silico* cfDNA Generation with Pleiades (a)** Distribution of generated-fragment insert sizes for Pleiades 600M versus ground truth; dashed lines mark the insert-size distribution peaks (mean peak distance indicated). **(b)** Longest common subsequence (LCS) length between generated and true fragments, expressed relative to the ground-truth length. **(c)** Methylated-cytosine context ratios: deviation from the expected proportion of methylated calls in each sequence context (CpG, CHH, CHG) for generated fragments (Evo 2 7B, Pleiades 600M, Pleiades 7B) relative to ground truth.

Supplementary Table S3 lists all the regions used in the analysis in section 2.3. The stable nucleosome was determined using NucPosDB[1]. To identify these regions, we selected 80 high-coverage 1kb intervals from our cfDNA test-set biosamples that do not overlap any repetitive elements (as defined by the UCSC Genome Browser's RepeatMasker track).

**Table S3**: 1kb Regions Evaluated for cfDNA Generation

| Chromosome | Region Start | Region End | Stable Chromatin |
|---|---|---|---|
| chr1 | 16,617,000 | 16,618,000 | No |
| chr1 | 107,143,000 | 107,144,000 | Yes |
| chr2 | 89,768,000 | 89,769,000 | No |
| chr2 | 174,338,000 | 174,339,000 | Yes |
| chr2 | 192,197,000 | 192,198,000 | Yes |

*Continued on next page...*

[1] https://generegulation.org/stable-nucleosomes/

Table S3 (continued)

| Chromosome | Region Start | Region End | Stable Nucleosome |
|---|---|---|---|
| chr3 | 107,527,000 | 107,528,000 | No |
| chr3 | 113,945,000 | 113,946,000 | Yes |
| chr4 | 105,149,000 | 105,150,000 | No |
| chr4 | 183,101,000 | 183,102,000 | Yes |
| chr5 | 120,466,000 | 120,467,000 | No |
| chr5 | 141,363,000 | 141,364,000 | Yes |
| chr5 | 146,379,000 | 146,380,000 | Yes |
| chr6 | 1,514,000 | 1,515,000 | No |
| chr6 | 10,885,000 | 10,886,000 | No |
| chr6 | 53,648,000 | 53,649,000 | No |
| chr6 | 56,543,000 | 56,544,000 | No |
| chr6 | 70,960,000 | 70,961,000 | No |
| chr6 | 79,079,000 | 79,080,000 | No |
| chr6 | 84,770,000 | 84,771,000 | No |
| chr6 | 96,833,000 | 96,834,000 | Yes |
| chr6 | 98,838,000 | 98,839,000 | No |
| chr7 | 28,177,000 | 28,178,000 | Yes |
| chr7 | 28,850,000 | 28,851,000 | No |
| chr7 | 29,196,000 | 29,197,000 | Yes |
| chr8 | 30,386,000 | 30,387,000 | No |
| chr8 | 64,584,000 | 64,585,000 | No |
| chr8 | 66,173,000 | 66,174,000 | No |
| chr8 | 66,922,000 | 66,923,000 | Yes |
| chr8 | 80,488,000 | 80,489,000 | No |
| chr8 | 123,271,000 | 123,272,000 | No |
| chr8 | 124,369,000 | 124,370,000 | No |
| chr9 | 968,000 | 969,000 | Yes |
| chr9 | 86,942,000 | 86,943,000 | Yes |
| chr9 | 89,003,000 | 89,004,000 | No |
| chr10 | 22,314,000 | 22,315,000 | No |
| chr10 | 133,526,000 | 133,527,000 | Yes |
| chr11 | 59,062,000 | 59,063,000 | Yes |
| chr11 | 125,168,000 | 125,169,000 | No |
| chr12 | 67,271,000 | 67,272,000 | Yes |
| chr12 | 79,687,000 | 79,688,000 | No |
| chr13 | 46,384,000 | 46,385,000 | Yes |
| chr13 | 99,976,000 | 99,977,000 | No |

*Continued on next page...*

*Table S3 (continued)*

| Chromosome | Region Start | Region End | Stable Nucleosome |
|---|---|---|---|
| chr14 | 19,344,000 | 19,345,000 | No |
| chr14 | 61,697,000 | 61,698,000 | Yes |
| chr15 | 20,360,000 | 20,361,000 | No |
| chr15 | 35,118,000 | 35,119,000 | No |
| chr15 | 52,567,000 | 52,568,000 | No |
| chr15 | 66,295,000 | 66,296,000 | Yes |
| chr15 | 67,523,000 | 67,524,000 | No |
| chr15 | 81,004,000 | 81,005,000 | No |
| chr15 | 96,335,000 | 96,336,000 | No |
| chr16 | 46,390,000 | 46,391,000 | No |
| chr16 | 79,596,000 | 79,597,000 | Yes |
| chr17 | 44,828,000 | 44,829,000 | Yes |
| chr17 | 70,166,000 | 70,167,000 | No |
| chr18 | 3,497,000 | 3,498,000 | No |
| chr18 | 3,502,000 | 3,503,000 | No |
| chr18 | 31,103,000 | 31,104,000 | No |
| chr18 | 57,356,000 | 57,357,000 | Yes |
| chr18 | 70,204,000 | 70,205,000 | No |
| chr18 | 80,146,000 | 80,147,000 | No |
| chr19 | 266,000 | 267,000 | No |
| chr20 | 34,214,000 | 34,215,000 | No |
| chr20 | 59,943,000 | 59,944,000 | Yes |
| chr21 | 10,625,000 | 10,626,000 | No |
| chr21 | 32,414,000 | 32,415,000 | Yes |
| chr22 | 11,624,000 | 11,625,000 | No |
| chr22 | 25,567,000 | 25,568,000 | Yes |

### S3: Cell Type-of-Origin

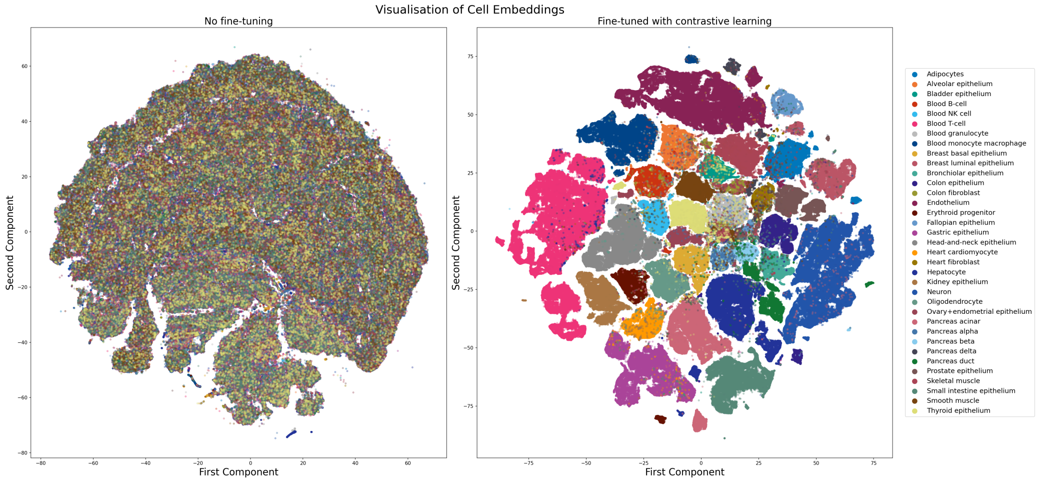


**Fig. S4**: Pleiades 90M Embeddings reduced to 2 dimensions using UMAP and colour-coded by cell type. The left is the pretrained model's representations with no discernible separation between cell types. The right image shows the model representations following fine-tuning.

| Mixture Name | B cell | T cell | Monocyte & Macrophage | Neuron | Oligodendrocyte | Hepatocyte |
|---|---|---|---|---|---|---|
| 1-Cell Mix | 1 | | | | | |
| 1-Cell Mix | | 1 | | | | |
| 1-Cell Mix | | | 1 | | | |
| 1-Cell Mix | | | | 1 | | |
| 1-Cell Mix | | | | | 1 | |
| 1-Cell Mix | | | | | | 1 |
| 2-Cell Mix | 0.5 | 0.5 | | | | |
| 2-Cell Mix | | | 0.5 | 0.5 | | |
| 2-Cell Mix | | 0.5 | | | | 0.5 |
| 3-Cell Mix | | 0.33 | | 0.34 | 0.33 | |
| 3-Cell Mix | | | | 0.34 | 0.33 | 0.33 |
| 3-Cell Mix | 0.33 | | | 0.34 | | 0.33 |
| 4-Cell Mix | 0.25 | 0.25 | 0.25 | 0.25 | | |
| 6-Cell Mix | 0.17 | 0.17 | 0.17 | 0.16 | 0.16 | 0.17 |

**Table S4**: Cell-Type mixture proportions used for deconvolution benchmarking. Blank entries indicate zero proportion.

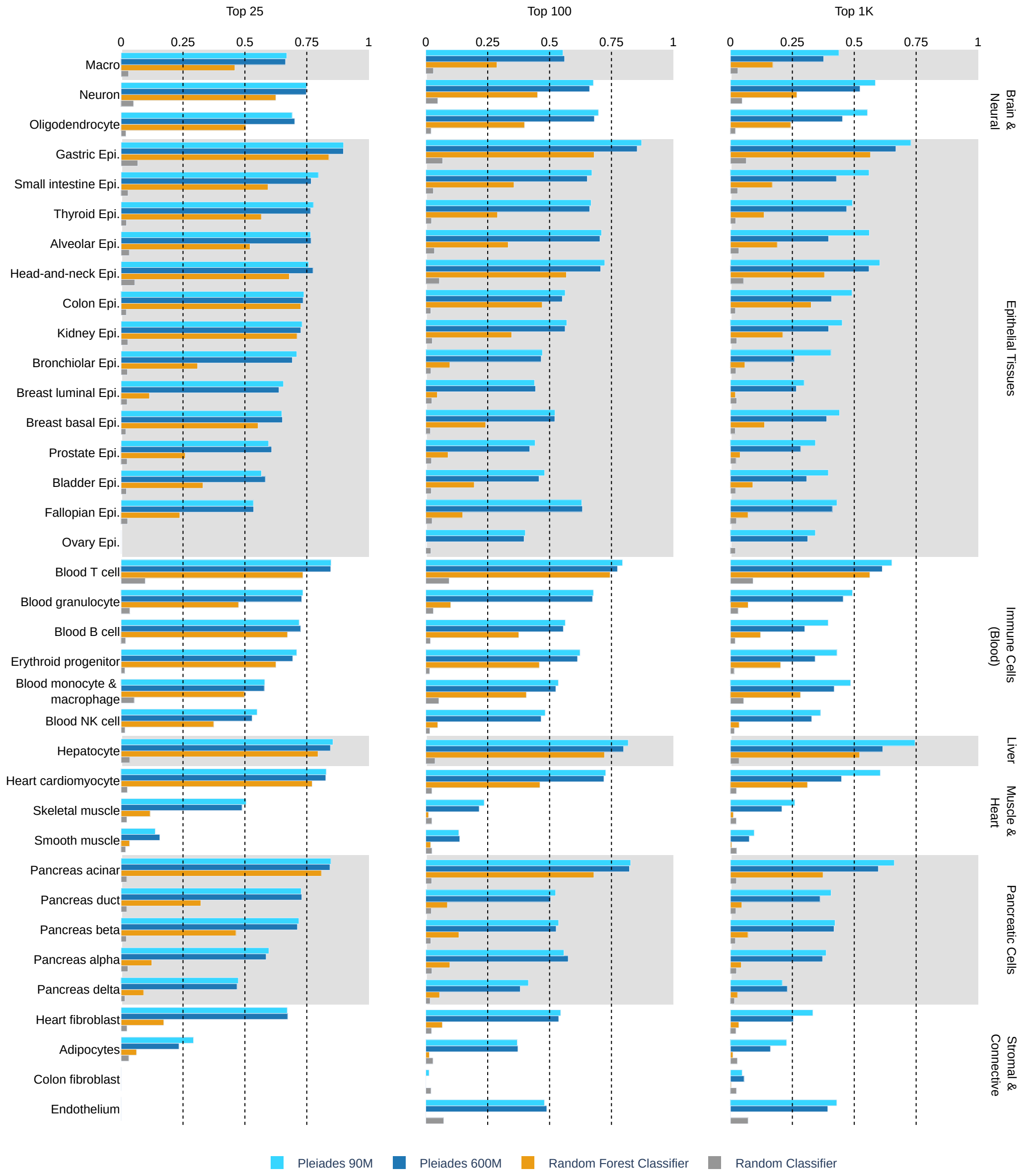


**Fig. S5**: CToO F1 scores over official Top 25, Top 100 and Top 1000 markers called from training data.

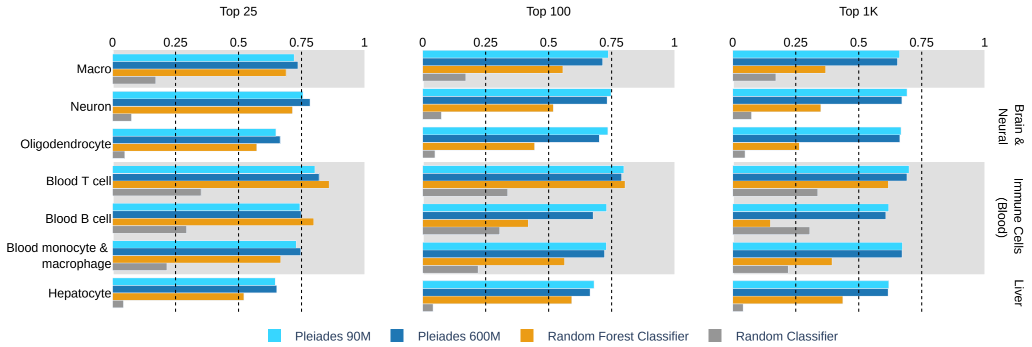


**Fig. S6**: Out-of-distribution CToO F1 scores over official Top 25, Top 100 and Top 1000 markers called from training data.

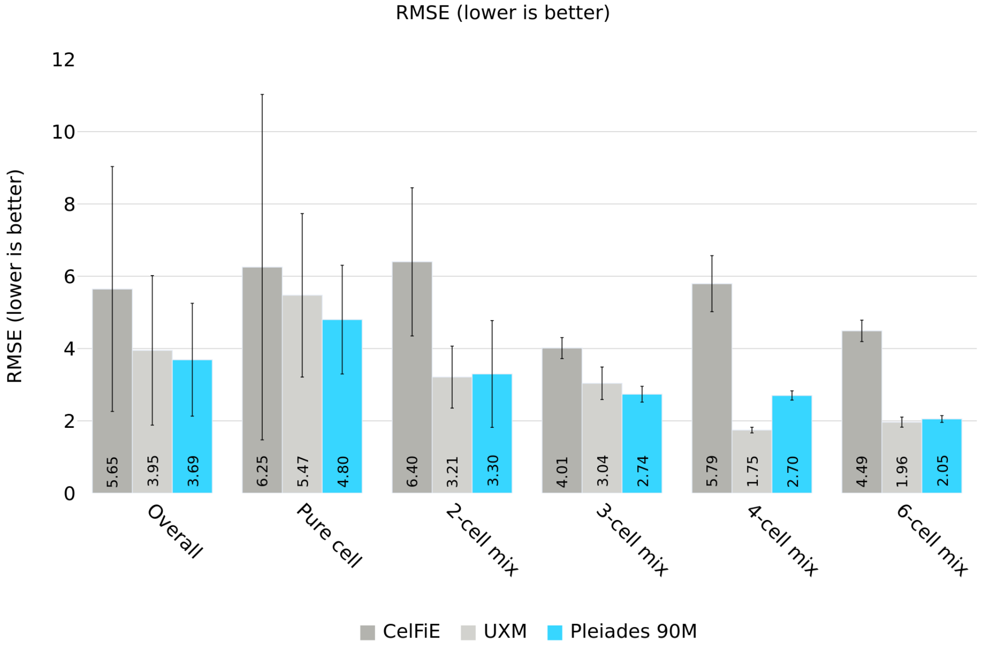


**Fig. S7**: For the deconvolution benchmark shown in Fig. 2e, we additionally report the root mean squared error (RMSE). Bars indicate the mean RMSE between true cell type ratio and estimated ratios (lower is better); error bars indicate the standard deviation.

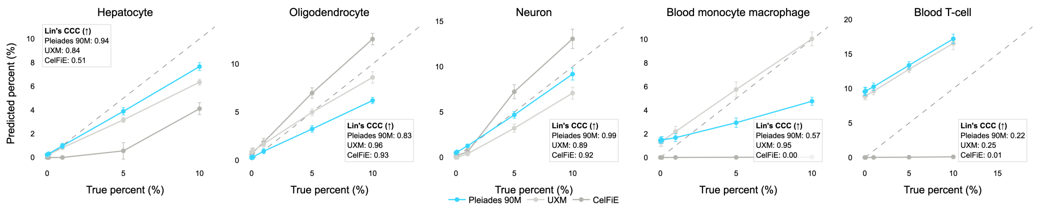


**Fig. S8**: **Additional *in silico* deconvolution experiments.** Predicted versus true cell type proportion for variable-proportion *in silico* spike-in experiments. Starting from a pool of B cell fragments, each of five cell types (hepatocyte, oligodendrocyte, neuron, monocyte/macrophage, T cell) was spiked in at 0%, 0.1%, 1%, 5%, and 10% frequency. Points show the mean predicted proportion across 5 random seeds; error bars indicate standard deviation. The dashed line indicates perfect concordance (y = x). Pearson correlation coefficient (r), Lin's concordance correlation coefficient (CCC), and RMSE are reported for each method per cell type.

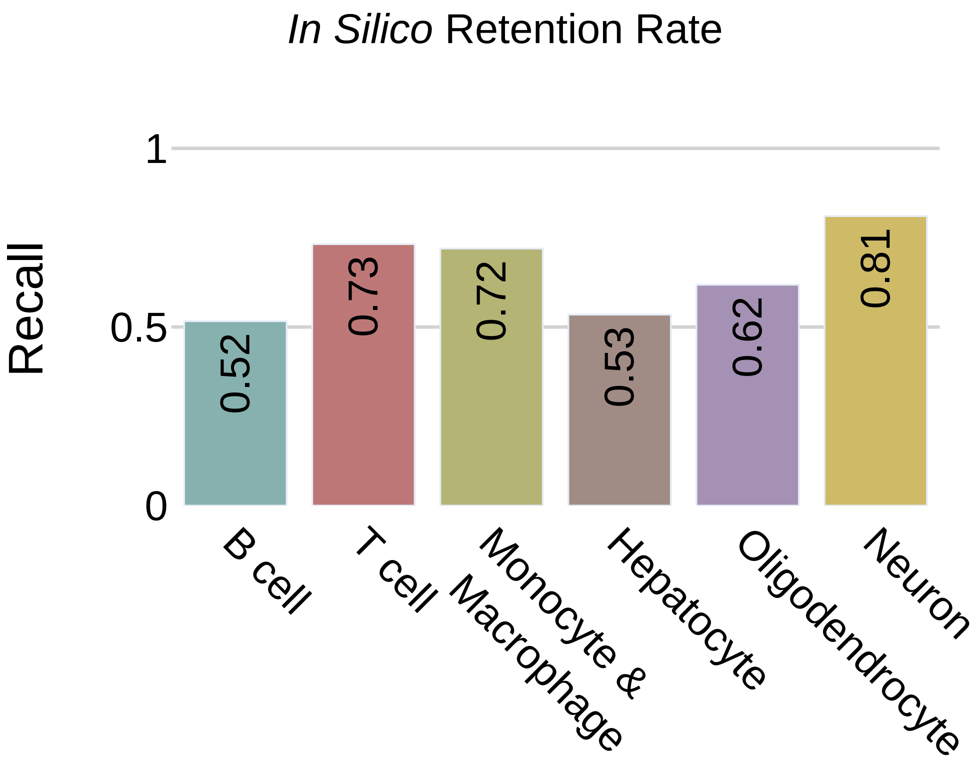


**Fig. S9**: Out-of-distribution CToO Recall over Top 1000 markers called from training data.

### S4: Neurodegenerative Disease Diagnosis

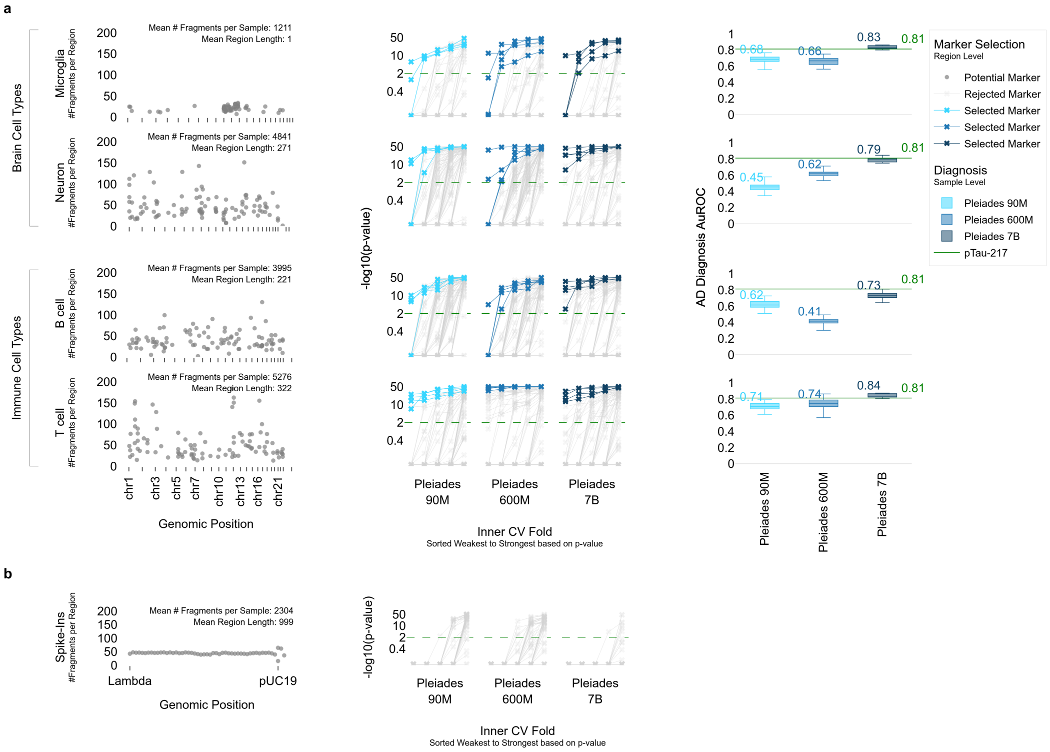


**Fig. S10**: **AD Marker Discovery Process with Pleiades (a)** AD marker discovery pipeline on an example fold of the outer five-fold split. The train set is divided into five inner folds. Starting from broad genomic regions across four cell type marker region sets, we select the regions with AUROC $> 0.6$ with $p$-value $< 0.01$ for at least four out of five inner folds. If more than four regions pass this test, we sort by number of folds passing the test and average AUROC, both descending, and select only top four. If any regions are selected, we train the region-level and sample-level models on those selected regions only. Final sample level performance in this example outer fold is shown along with the starting regions and the T-test outcomes for region selection. **(b)** Same method applied to pUC19 and Lambda DNA spike-ins; no signal was detected and the model rejected all regions.

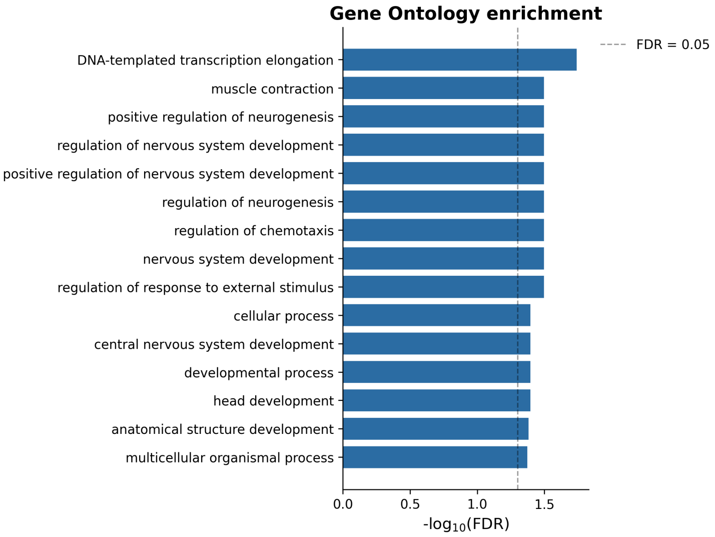


**Fig. S11**: **GO term enrichment in HAT-selected marker regions.** Horizontal bars show $-\log_{10}(\mathrm{FDR})$ from GREAT binomial test for the top enriched GO terms. Analysis was performed on all unique top-4 regions pooled across all cell types (neuron, microglia, B cell, T cell), tested against the starting candidate DMRs as background. Blue bars indicate FDR $< 0.05$; dashed line marks the FDR $= 0.05$ threshold. Of 724 GO:BP terms tested, 18 were significantly enriched (FDR $< 0.05$), with prominent representation of neurodevelopmental processes including nervous system development, regulation of neurogenesis, and brain development.

| Proteomic Marker | Test Direction | p−value (raw) | p−value (BH adjusted) |
|---|---|---|---|
| AB40 (pg/ml) | Greater | 0.0001 | 0.0005 |
| AB42 (pg/ml) | Greater | 0.0000 | 0.0001 |
| $\frac{\text{AB42 (pg/ml)}}{\text{AB40 (pg/ml)}}$ | Greater | 0.0010 | 0.0026 |
| NfL (pg/ml) | Greater | 0.0069 | 0.0139 |
| GFAP (pg/ml) | Greater | 0.0126 | 0.0168 |
| pTau-181 (pg/ml) | Greater | 0.0105 | 0.0168 |
| pTau-217 (pg/ml) | Greater | 0.0263 | 0.0301 |
| pTau-217 (pg/ml) | Less | 0.9737 | 0.9737 |

**Table S5**: Comparison of Pleiades 7B AUROC (average pooled on all cell type markers) vs. Proteomic Markers using one-sided paired t-test. Test direction column represents the direction used during testing, with *Greater* meaning Pleiades AUROC $>$ proteomic AUROC and *Less* meaning Pleiades AUROC $<$ proteomic AUROC. For all markers other than pTau-217, only *Greater* direction was used due to the large distance in means in favour of Pleiades 7B.

### S5: Mechanistic Interpretability of Pleiades AD Classification

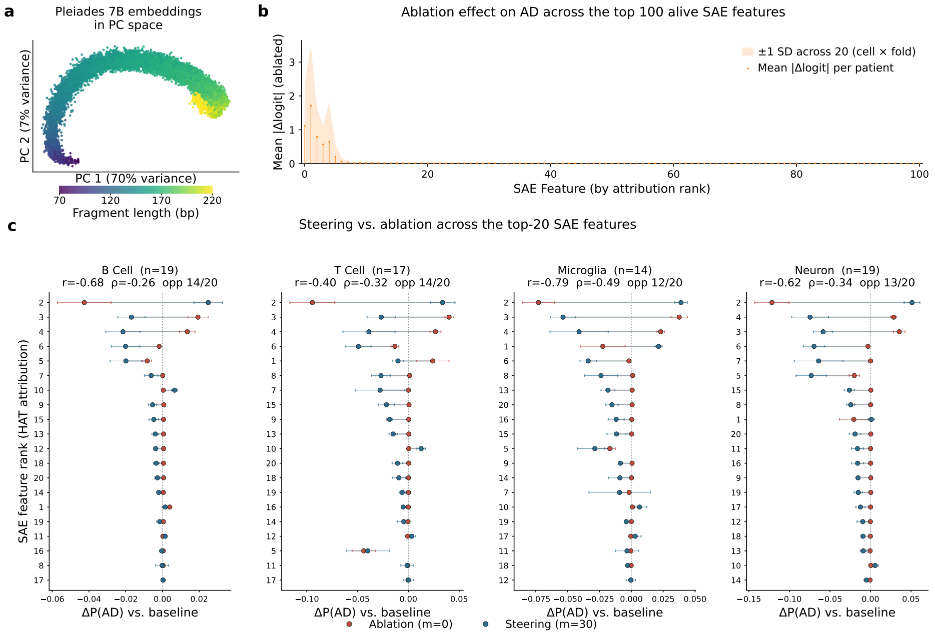


**Fig. S12**: **(a)** PCA manifold of post-trained Pleiades 7B CLS-token activations, coloured by fragment length. Two-dimensional scatter of the first two principal components of CLS-token activations from the post-trained Pleiades 7B checkpoint on the AD cohort, restricted to the top-4 regions per cell type. Each point is one fragment; colour encodes fragment length. Fragment length explains 49.66% of the variance, with PC1 capturing 70.50% and PC2 6.78%. **(b)** Ablation effect on AD prediction across the top 100 alive SAE features. Each alive SAE feature was ablated globally; its decoder contribution subtracted from every fragment's CLS activation (steering multiplier $=$ 0, no region mask). The resulting change in the HAT model's AD log-odds was measured on every held-out test patient. Points show the mean per-patient —$\Delta$logit— for each feature, averaged over the 20 (cell type $\times$ fold) experiments; the shaded band is $\pm 1$ SD across those experiments. Features are ordered along the x-axis by descending mean attribution; the 100 highest-ranked of the alive features are shown. The ablation effect concentrates in the few top-attributed features and decays to near zero for the remainder, indicating that AD-relevant signal is carried by a small subset of features. **(c)** Steering versus ablation across the top-20 SAE features, per cell type. For each of the top-20 attributed SAE features, the change in predicted AD probability, $\Delta$P(AD), relative to baseline (multiplier $=$ 1) is shown for two opposite interventions: feature ablation (red, multiplier $=$ 0) and feature steering at the strongest amplification (blue, multiplier $=$ 30). Each dumbbell connects a feature's mean ablation and steering $\Delta$P(AD) across unique genomic regions (pooled across folds by region identity); horizontal bars are the SEM across those regions, and rows are ordered by dumbbell span (largest at top). Panels show the four cell types (B cell, T cell, microglia, neuron); n is the number of unique regions. Annotations report the Pearson (r) and Spearman ($\rho$) correlation between the ablation and steering effects across the 20 features, and the number of sign-opposite features (those for which amplifying and removing the feature move P(AD) in opposite directions) — the expected signature of causally consistent features.

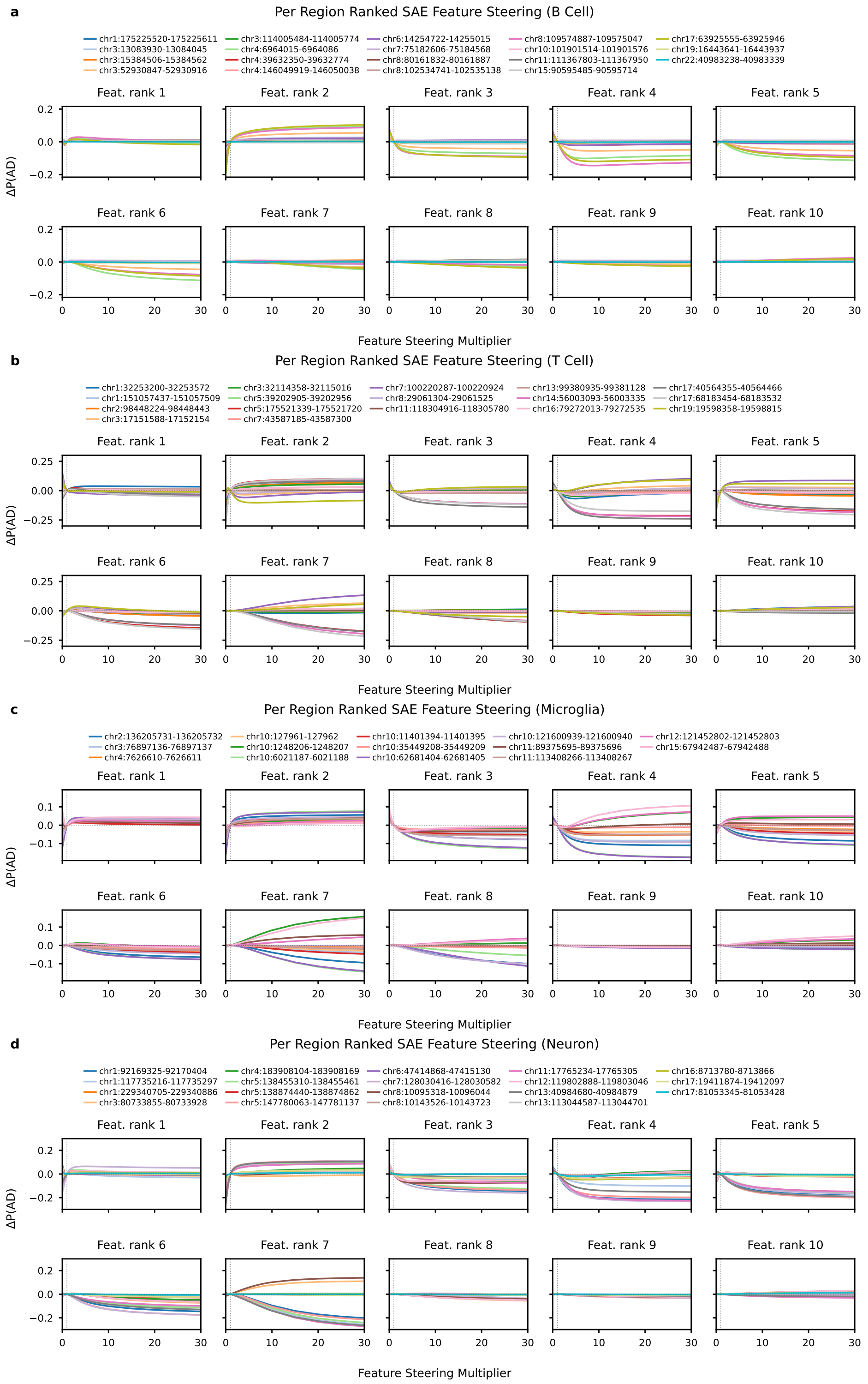


**Fig. S13**: **Per region steering response curves for the top-10 SAE features by cell type. (a–d)** One $2 \times 5$ panel grid per cell type (B cell, T cell, microglia, neuron), one subplot per attribution-ranked SAE feature. Curves show $\Delta P(\mathrm{AD})$ versus the steering multiplier $m$ (14-point grid, half-steps in $[0, 3]$ and unit steps to 10; $m = 1$ is the unsteered baseline). Lines are coloured by region identity (14–19 unique regions per cell type). Features are indexed by rank position; curves are averaged over patients within a fold and then over folds in which the region appears.

| **Comparison** | $t$ | $p$ (Dunnett) | **sig** |
|---|---|---|---|
| **B Cell** | | RM-ANOVA $F(3,12) = 9.00$, $p = 0.0021$ (**) | |
| Frag. Len. (rand.) vs. Pleiades 7B | -5.01 | 0.0009 | *** |
| Frag. Len. (DELFI) vs. Pleiades 7B | -3.27 | 0.0174 | * |
| Frag. Len. (Pleiades) vs. Pleiades 7B | -1.91 | 0.1876 | ns |
| **T Cell** | | RM-ANOVA $F(3,12) = 14.17$, $p = 0.0003$ (***) | |
| Frag. Len. (rand.) vs. Pleiades 7B | -5.28 | 0.0006 | *** |
| Frag. Len. (DELFI) vs. Pleiades 7B | -4.41 | 0.0022 | ** |
| Frag. Len. (Pleiades) vs. Pleiades 7B | -0.59 | 0.8861 | ns |
| **Microglia** | | RM-ANOVA $F(3,12) = 4.48$, $p = 0.0249$ (*) | |
| Frag. Len. (rand.) vs. Pleiades 7B | -3.27 | 0.0176 | * |
| Frag. Len. (DELFI) vs. Pleiades 7B | -3.06 | 0.0253 | * |
| Frag. Len. (Pleiades) vs. Pleiades 7B | -2.31 | 0.0955 | ns |
| **Neuron** | | RM-ANOVA $F(3,12) = 5.11$, $p = 0.0165$ (*) | |
| Frag. Len. (rand.) vs. Pleiades 7B | -3.41 | 0.0136 | * |
| Frag. Len. (DELFI) vs. Pleiades 7B | -2.76 | 0.0432 | * |
| Frag. Len. (Pleiades) vs. Pleiades 7B | -0.84 | 0.7405 | ns |

**Table S6**: Per-cell-type one-way repeated-measures ANOVA followed by Dunnett's post-hoc test of held-out 5-fold cross-validated AD-classification $AUROC$ across the four classifiers: *Frag. Len. (rand.)*, a linear fragment-length model on the median fragment length of four random regions (averaged over 20 random four-region draws per fold); *Frag. Len. (DELFI)*, the published DELFI baseline at 151 bp; *Frag. Len. (Pleiades)*, a linear fragment-length model on the median fragment length of the four attribution-selected Pleiades regions per fold; and *Pleiades 7B*, the sample-level classifier. Within each cell type, fold is the repeated factor ($n = 5$ folds): a one-way repeated-measures ANOVA across the four models (group header row, $F(3,12)$) tests the omnibus model-type effect, and Dunnett's post-hoc test then contrasts each fragment-length model against the *Pleiades 7B* control, with the reported two-sided $p$-values family-wise-error-controlled across the three control-vs-treatment comparisons within that cell type. The `sig` column reports the significance marker (*** $p < 0.001$, ** $p < 0.01$, * $p < 0.05$, ns otherwise), applied to the ANOVA $p$ in the header rows and to the Dunnett $p$ in the contrast rows.
